# Bridging Ecological Inference and Decision Optimization for Conservation Using Artificial Intelligence

**DOI:** 10.64898/2026.08.13.744541

**Authors:** Hyun Seok Yoon, Charles B. Yackulic, Abigail J. Lawson, Casey Wagnon, Kasey Pregler

**Affiliations:** Department of Fish, Wildlife, & Conservation Ecology, New Mexico State University, Las Cruces, NM; U.S. Geological Survey, Southwest Biological Science Center, Grand Canyon Monitoring and Research Center, Flagstaff, AZ; U.S. Geological Survey, New Mexico Cooperative Fish and Wildlife Research Unit, Las Cruces, NM

**Author notes:** Classification: Ecology; Environmental Sciences.

**Keywords:** Stochastic Dynamic Programming, Hierarchical Bayesian Modeling, Adaptive Management, Conservation Decision Science, Population Viability Analysis, Markov Decision Process

## Abstract

The ability to model the complex and uncertain population dynamics of endangered species has improved dramatically in recent decades. However, approaches to identify optimal decisions often require a simplified representation of population dynamics. This leads to a conundrum where managers may be unsure about the output of dynamic decision models because they rely on simplified assumptions of the underlying population dynamics. Here, by pairing integrated population models (IPM) that synthesize diverse ecological data with deep reinforcement learning (DRL) capable of optimizing decisions with high-dimensional uncertainty, we introduce a framework that delivers data-driven and ecologically detailed adaptive management strategies. We demonstrate its utility through application to the supplementation program for the endangered Rio Grande silvery minnow. Using our IPM-DRL framework, we developed an adaptive decision model that selects production and distribution decisions of the supplementation program in response to the observed demographic, hydrological, and genetic environment. The decision model outperformed all heuristic approaches in the simulation across management objectives that weighed persistence and effective population size-related genetic impact differently. For example, the currently deployed supplementation strategy performed 5.3% worse than the decision model under the persistence-focused objective scoring and 185% worse under the genetics-focused one. Analysis of the model’s decisions in relation to demographic and environmental covariates revealed that minimum sub-population size and total population size were primary drivers of the model’s decisions. The results demonstrate that the IPM–DRL framework offers a high-performing and interpretable decision-support tool for managing endangered species.

**Significance:** Conservation problems, like imperiled species management, are often challenging because the system dynamics are complex and uncertain. We demonstrate how combining an integrated population model that infers key demographic processes from noisy ecological data with a deep reinforcement learning framework that optimizes management actions addresses these challenges by generating high-performing supplementation strategies for a conservation-dependent species. Our approach embeds two decades of monitoring data within a multi-objective decision-making environment that accounts for ecological uncertainty. The result is a generalizable framework that links ecological inference directly to actionable policy outcomes, enabling scientists and managers to move beyond describing system states and processes toward identifying optimal management actions.

## Introduction

Endangered species management is a central component of biodiversity conservation (1). While all conservation problems navigate complex and uncertain socio-ecological dynamics (2), management planning for endangered species often necessitates a more detailed representation of the problem when permitted by available science for several reasons (3). First, legal statutes regarding the protection of populations and habitats of endangered species, combined with the ethical weight of preventing extinction, require careful consideration of the potential outcomes of management actions (4,5). Second, for imperiled species, there often exists detailed expert knowledge of species’ life-history traits and the environmental stressors driving populations toward a critical level (6). Though species and experts may prefer to capture the population dynamics and uncertainty in substantial detail when developing models for management planning (3), added complexity could come at the cost of understanding and transparency, reducing the likelihood of adoption (3,7). This further perpetuates the research-implementation gap pervasive in conservation science (8,9). However, the trade-off of making the decision model more complex is that it leaves little room for simplification, making it harder to solve for an optimal management solution.

Management of endangered species happens in dynamic and stochastic environments and under imperfect knowledge about system dynamics (10). Statistical population models commonly used to inform species management are often complex and explicitly characterize uncertainty across numerous parameters through hierarchical structures (11,12). For example, ecologists are increasingly developing integrated population models (IPMs) to synthesize diverse data sources (e.g., counts, mark–recapture, telemetry, reproduction) within a hierarchical framework to estimate posterior distributions of relevant demographic parameters (13,14). However, traditional methods for modeling optimal solutions become computationally infeasible as representations of population dynamics become more complex (15,16). Therefore, most stochastic dynamic decision models in conservation have heavily simplified the conservation problem to remain tractable (17–19).

Recent advancements in deep reinforcement learning (DRL) offer a path forward to approximating optimal decisions for complex stochastic dynamic decision problems. In a DRL modelling approach, an agent, or a virtual manager, chooses management actions and observes the response from the complex representation of system dynamics. By rewarding the agent based on positive outcomes, the neural network that represents the agent’s decision under different conditions converges over the course of a simulation. DRL models are widely applied in industries to tackle complex tasks (20,21), but there have been only a few applications of DRL in conservation (22,23). Given the complexity of population models and increasingly sophisticated ways to incorporate uncertainty in model simulations, DRL approaches represent a timely advancement to ensure that our nuanced ecological understanding informs management strategies for imperiled species.

Here, we show how combining population inference and decision optimization can improve endangered species management under uncertainty. We demonstrate the utility of the IPM–DRL framework for developing a management strategy for endangered species by applying it to the hatchery supplementation of the Rio Grande silvery minnow (*Hybognathus amarus*; RGSM).

The RGSM is a freshwater fish that has been extirpated from 95% of its historical range and is now restricted to the middle Rio Grande region (24). It is a short-lived species, with most individuals reproducing once, and abundance can fluctuate by orders of magnitude between years. This volatility elevates extinction risk during periods of poor recruitment or survival, requiring repeated management decisions. Its endangered status under the Endangered Species Act and the severely strained water resource in this arid region have made the conservation of RGSM both challenging and contentious (25–27). Since 2002, the U.S. Fish and Wildlife Service has led a captive breeding program to supplement the wild population. The program has been instrumental in preventing extinction, with broodstock management explicitly designed around the species’ reproductive biology to preserve genetic diversity (28,29). Due to sustained management intervention, extensive demographic and genetic data have been collected for RGSM through long-term monitoring, although the resulting data are highly heterogeneous (30,31), comprising different sources of monitoring activities (regular monitoring vs samples collected during other management activities), sampled habitat types (backwater pools vs runs), and changes in the number of sampling sites and the distribution of habitat types over time (30).

The supplementation program faces two major decisions annually: 1) how much fish to produce at the hatchery in the spring, and 2) how to distribute the produced fish in the fall across the three river-segment reaches, Angostura, Isleta, and San Acacia. The two main objectives of the supplementation program are to minimize the extinction risk of the population while also maintaining the genetic health of the population (28,29). While supplementation can affect population genetics through many mechanisms, in this paper, we focus on its effect on the inbreeding effective population size, hereafter referred to as effective population size. A population genetics model specific to RGSM does not exist, so we developed a model based on established population genetics theory and combine it with a previously parameterized population dynamics model. We compare the performance of dynamic supplementation models derived from the IPM–DRL framework, measured as average cumulative objective scores (rewards) across thousands of 50-year simulations, with other heuristic strategies, including the current strategy, designed around the trade-off that can exist between these dual objectives. The current strategy adjusts hatchery production inversely with spring-flow forecasts and allocates stocked fish to reaches where expected catch per unit effort is below one (SI1, Appendix A). In addition, we demonstrate how the framework can support other common conservation planning analyses, such as value-of-information assessments (32) and evaluations of alternative management plans. Through this analysis, we present DRL as a reliable tool for deriving practical and effective endangered species management and assessing new monitoring and management plans when coupled with other robust statistical tools.

## Results

### Using the IPM–DRL framework in endangered species management

The IPM–DRL framework links demographic inference from an IPM directly to a trained DRL decision model (Fig. 1). First, the IPM is used to infer key latent demographic states, such as reach– and age-specific population sizes, from heterogeneous monitoring data (Fig. 1a). These inferred demographic states, together with environmental covariates available to managers, provide the information used to generate management recommendations (i.e., production and distribution decisions for supplementation) from the DRL model. To ensure that the DRL model is trained under realistic ecological uncertainty, the posterior distributions of demographic process parameters (e.g., recruitment rate, mortality) from the IPM are used to propagate the uncertainty within the training simulation. The DRL model can be retrained periodically using parameter distributions from the refitted IPM as new monitoring data accrue, implementing an adaptive management cycle (Figure 1b) (33,34).

**Figure 1.**
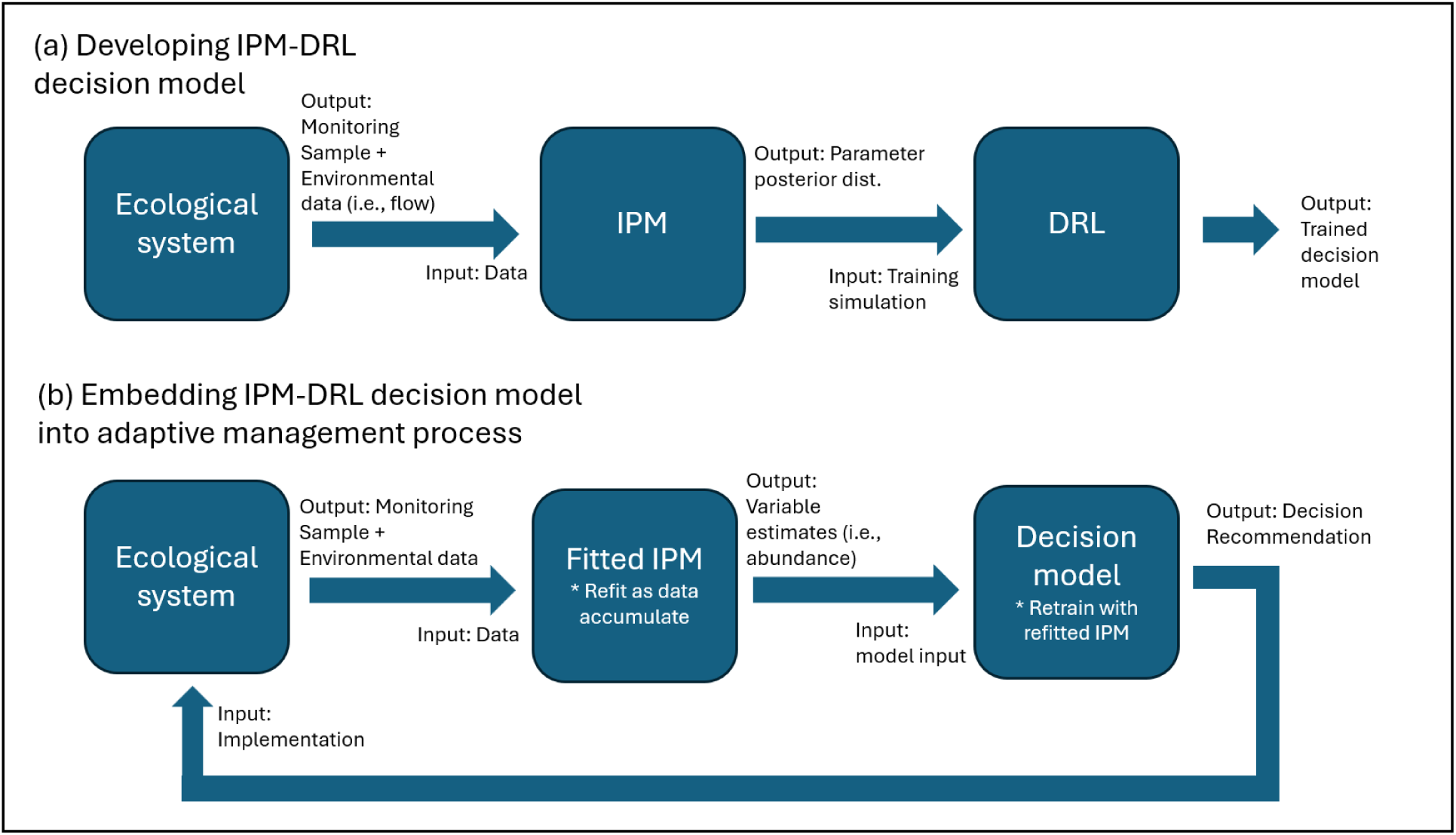
Workflow of the IPM–DRL framework for endangered species management. (a) Monitoring and environmental data are used to fit an integrated population model (IPM), and the resulting posterior parameter distributions are used to train a deep reinforcement learning (DRL) decision model through simulation. (b) Once deployed, the IPM uses new data to estimate system states (e.g., abundance), which are passed to the decision model to generate management recommendations. Implemented decisions then affect the ecological system, creating an adaptive management cycle as new data accumulate.

### Performance of supplementation decision models derived from DRL

The DRL algorithm adjusts both production and distribution decisions to maximize cumulative rewards over the simulation horizon. In RGSM supplementation, the reward reflects its dual objectives of persistence and genetic impact. Specifically, it is a sum of the annual value of persistence of species and the genetic impact of stocking that year. The value of persistence represents the relative importance a manager places on species persistence in a given year relative to the genetic impact incurred from stocking in that year. Genetic impact is quantified as the difference in log effective population size between post-stocking conditions and a counterfactual scenario without stocking. Effective population size is tracked independently of the demographic processes governing persistence. A larger effective population size signifies lower rate of inbreeding, which helps preserve the genetic diversity, especially in terms of heterozygosity. The effective population size decreases when the hatchery effective population size is small compared to the wild effective population size and there is a large contribution of hatchery broodstock to the next generation (SI1, Appendix A). Increases in effective population size from supplementation is also possible if the wild effective population size is sufficiently small. However, a larger amount of stocking also increases the probability of persistence, leading to a potential trade-off between genetic diversity and persistence in some years under intermediate or high wild effective population size.

Consequently, decision models trained with higher weight on persistence in the reward function (Eq 1.) prioritize persistence over genetic diversity, yielding outcomes with higher persistence probabilities but lower long-term effective population sizes (Fig. 2a). The Pareto curve produced by training the DRL model on various persistence values allows the manager to select the model that produces the outcome that matches their preference in the persistence-genetics trade-off. The curve illustrates that DRL model can produce higher persistence than any other heuristics, thus preferable even with a high weight on persistence objective. We note that the long-term effective population sizes under all strategies in Fig. 2a, including the current strategy, remain sufficiently large to avoid substantial loss of genetic diversity, reflecting the overall effective current genetic management strategy for the supplementation. Nevertheless, the DRL model’s clear advantage over heuristic strategies in terms of genetic impact (Fig. 2a) highlights the value of our framework in systems where genetic risks are a larger concern (e.g., salmonids) (35).

**Figure 2.**
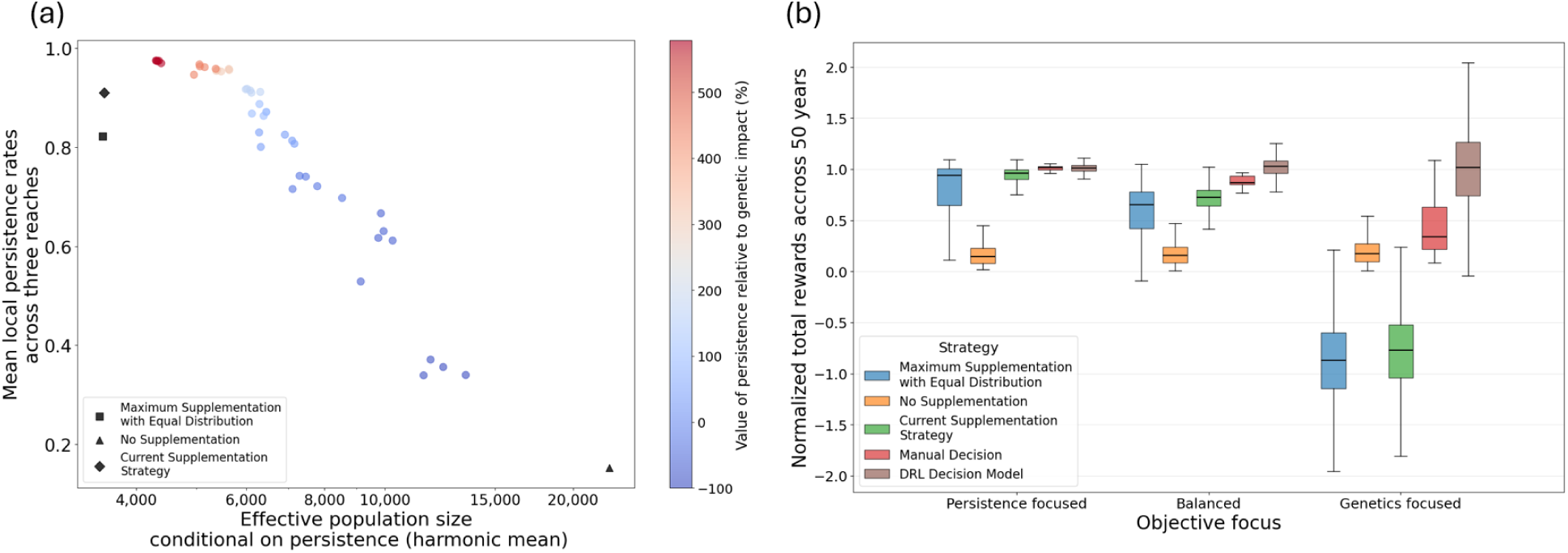
(a) Effective population size conditional on persistence (harmonic mean in log_10_ scale) and local persistence rate in a simulated 50-year supplementation program with varying value of persistence. (b) Average total rewards received from a 50-year supplementation using different heuristic strategies under persistence focused (value of persistence = 589%), balanced (172%), and genetics focused (−32%) objectives (n=20 for manual decision, n=3,000 for all others). The rewards are normalized by the average total rewards received from the DRL-derived decision model. Value of persistence is measured in percentages to the average annual genetic impact when following the current supplementation strategy. Local rate of persistence was defined as the number of years the local population wasn’t extinct in a 50-year period, averaged across three reaches.

DRL-derived decision models outperformed all heuristic supplementation strategies considered — no supplementation, the current management strategy, maximum supplementation with equal distribution among reaches, and a manually specified decision rule — under all weightings of persistence over genetics objectives (Fig. 2). Under no supplementation, local persistence rates were very low (Fig. 2a), resulting in substantially lower median total rewards than the DRL decision model under both the persistence-focused and genetics-focused objectives (Fig. 2b).

Conversely, the manual, current, and maximum supplementation strategies emphasized persistence over genetic impact, achieving higher local persistence rates but lower long-term effective population sizes (Fig. 2a). Their median rewards were similar to those of the DRL model under the persistence-focused objective but substantially lower under the genetics-focused objective, with some strategies producing negative median rewards (Fig. 2b). Both the current and manual strategies struggled to navigate population genetic dynamics and time the stocking in a way that reduce negative genetic impact.

Decisions concerning production and stocking distribution from DRL models showed a distinct contrast across gradients of value of persistence (Fig. 3). Average production level, defined as the proportion of the hatchery production capacity utilized across fifty years of supplementation, decreased with lower value of persistence to reduce the impact of stocking on genetic diversity (Fig. 3a). The year-to-year production was mostly binary, where the manager either produced the maximum amount or none (Fig. 3c). Years with lower population size or lower minimum local population tended to trigger maximum production, and the threshold line for trigger moved up with higher value of persistence (SI2, Fig. S3). Production decisions were also strongly conditioned on spring flow forecasts (SI2, Fig. S4,5).

**Figure 3.**
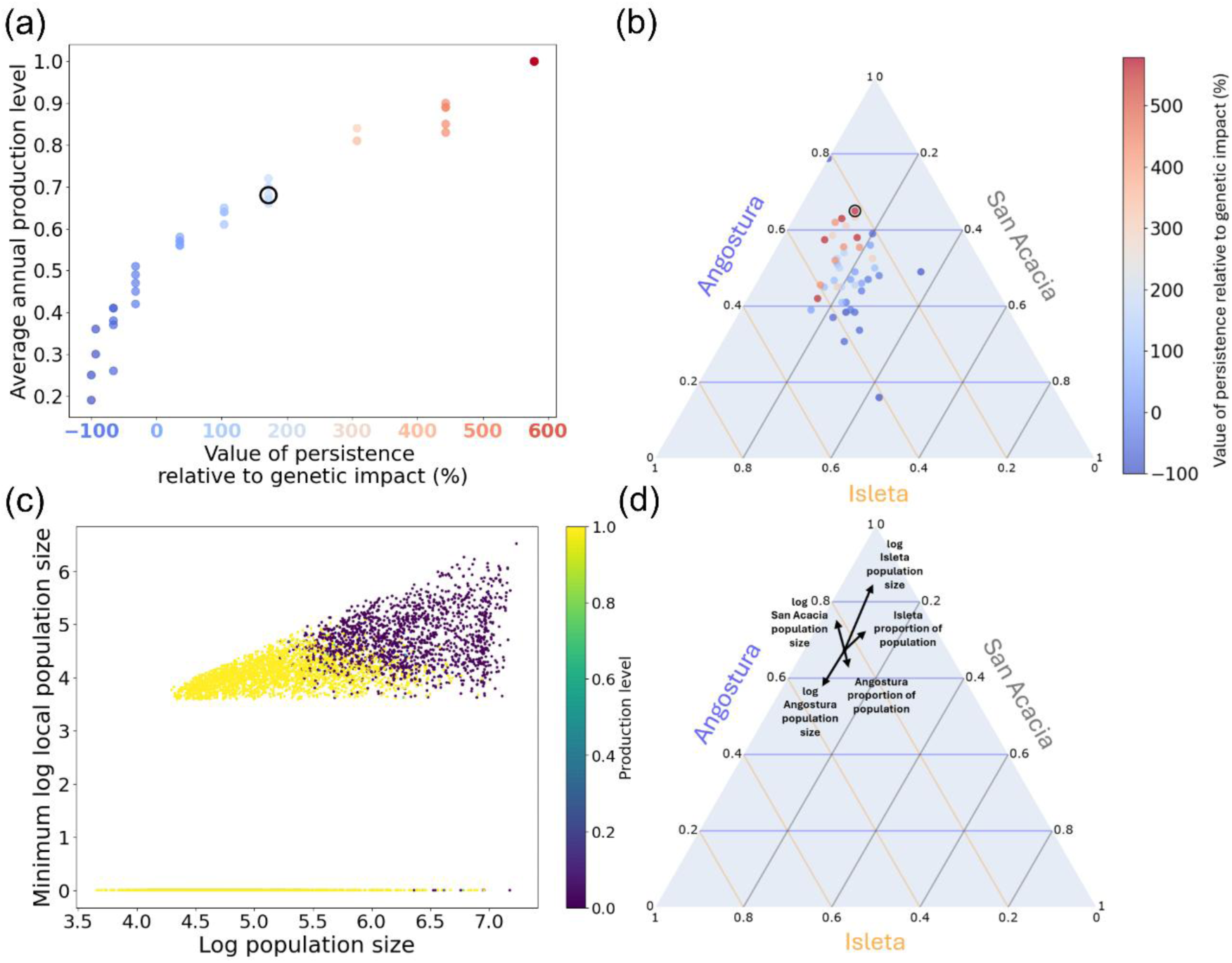
(a) Average production level and (b) stocking distribution across the three river reaches (triangle edges) in a DRL-derived decision model across a 50-year supplementation program; and (c) annual production level decision described by log_10_ of minimum local population size and log_10_ of total population size under the balanced objective and (d) the effect of changes of five most influential covariates that describes the stocking distribution across the reaches under the persistence-focused objective. Production level of 1 equates to producing the maximum capacity of the hatchery, which is 200,000 fish. Effect of covariates were measured by their changes by one standard deviation. The origin of arrows is the stocking distribution predicted by the median value of covariates from Dirichlet regression on the stocking decisions. Annual decisions (c, d) are sampled across a hundred 50-year simulations. The circled models in (a) and (b) indicate the ones used for creating (c) and (d), respectively.

Mean stocking distribution with higher persistence values consistently emphasized stocking in the Angostura reach with 50 to 70% of production, followed by 20 to 40% in Isleta, and then 0 to 20% in San Acacia (Fig. 3b). Emphasis on Angostura was mostly to compensate for the higher average mortality rate in the Angostura reach (SI1, Table S1). As the value of persistence decreased, DRL models no longer converged on a single dominant stocking pattern, indicating that the precise allocation of fish across reaches became less influential than the production decision, which affects genetic diversity more strongly. In year-to-year stocking decisions, a higher local population or proportion of the total population in a reach typically led to less stocking in that reach to allocate more elsewhere (Fig. 3d).

### Evaluating management flexibility and information value

Beyond identifying optimal supplementation strategies under the baseline management scenario, the IPM–DRL framework enables analyses that are common in conservation planning: quantifying the value of improved information and evaluating the benefits of increased management flexibility.

Increased management flexibility through the ability to carry-over hatchery fish to the following year increased performance only under the persistence-focused objective by 1.9% on average.

The DRL model leveraged this flexibility to reserve fish produced in high-abundance years for potentially poor conditions in the subsequent year (Fig. 4). In low abundance years, the model either did not carry-over if few fish had been retained from previous year (Fig. 4a), or carry-over at a limited rate if there were enough fish retained from last year (Fig. 4b), indicating a strategic use of carry-over to buffer against population declines. Under balanced (persistence 172% more important) or genetics-focused objectives, however, carry-over provided little benefit since the decision model already produced less than quarter of the fifty years under the baseline scenario, making the added flexibility largely unnecessary.

**Figure 4.**
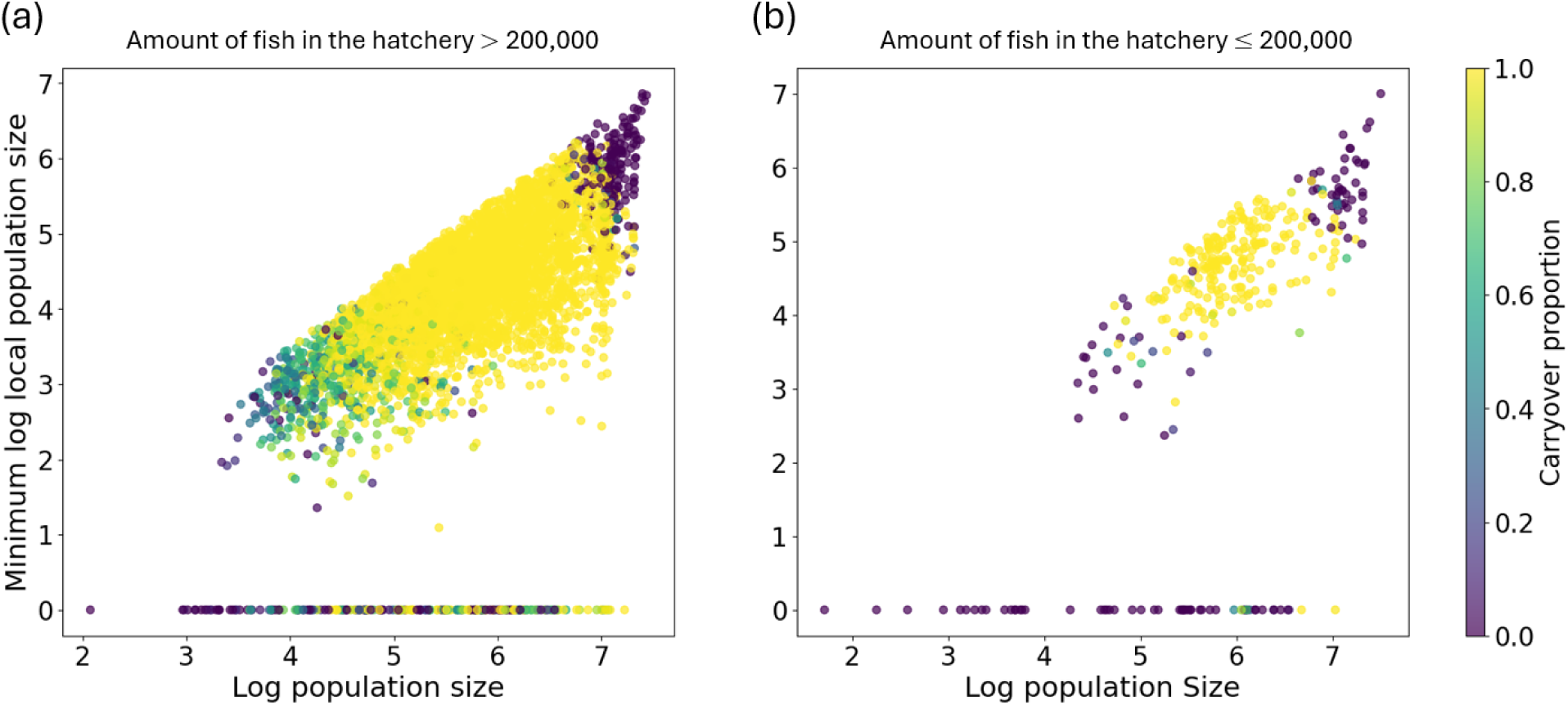
Annual carry-over decisions from a DRL-derived decision model under persistence-focus objective plotted over log_10_ of minimum local population size and log_10_ of total population size when the amount of fish in the hatchery is (a) greater than 200,000 or less than (b) 200,000. Carry-over proportion of 1 indicates deferring the stocking of all the fish produced that year to the next year. The decisions are sampled across a hundred 50-year simulations.

IPM–DRL framework is also conducive to assessing the expected value of perfect information (32) because the uncertainty around the population parameters is clearly defined by the posterior distribution from the IPM. We measured the expected value of perfect information when decision models were trained under scenarios where either winter mortality or reach-specific carrying capacity parameters were observed perfectly. Under the perfect information on these parameters, the performance improved by less than 1% across all persistence values. This negligible gain is consistent with many values of information analyses in conservation, where the expected value of perfect information tends to be small (36), indicating that resolving the uncertainty in these demographic processes would yield only a minor improvement relative to the baseline scenario.

## Discussion

Quantitative models for guiding endangered species management are an integral part of evidence-based and systematic conservation (37). However, surveys and literature reviews indicate practitioners often do not adopt quantitative decision-support tools, one of the reasons being the perception that there are too many variables in the system for simplistic models to be useful (7,38). For instance, in the development of a resource allocation tool for species listed on the Endangered Species Act at the U.S. Fish and Wildlife Service, one of the main hurdles for the working group to overcome was the sentiment that quantitative approaches are only useful for simple decision context (3).

Here, we demonstrated that integrating inferential statistical models with artificial intelligence-based decision frameworks can provide a powerful approach for managing endangered species in complex, dynamic, and uncertain environments. In our application to the RGSM supplementation, the framework derived dynamic supplementation decision models that outperformed heuristic approaches across different objectives in our model. These decision models successfully navigated, under different management objective weights, the trade-off dynamics commonly encountered in species conservation, such as balancing genetic impact against population persistence (39) and allocating resources across spatially structured populations (40), in a high-dimensional continuous environmental space. In particular, DRL-derived strategies outperformed current and alternative supplementation approaches most strongly under the genetics-focused objectives by avoiding stocking in years when genetic impacts were expected to be highly negative. Under persistence-focused objectives, DRL models achieved better performance by identifying stocking distributions that maximized persistence across all three reaches, primarily by allocating a greater proportion of hatchery fish to the Angostura reach, where mortality rates are highest.

A common criticism of DRL and stochastic dynamic programming in general is their lack of interpretability (10). The black box nature of neural networks and other complex modeling frameworks can limit user trust and hinder adoption in applied management contexts (8,41). However, by analyzing the DRL model’s decisions with biologically relevant covariates, we demonstrate direct ecological interpretation of the learned model. For instance, the current hatchery production plan reduces production when forecasted spring flow is high, reflecting favorable conditions for RGSM, thereby lessening the need for excessive stocking that may exacerbate genetic impacts. However, too little stocking may not sufficiently counter future demographic or genetic losses. DRL models offer strategies to balance these tradeoffs for genetics-, persistence-focused or balanced objectives to inform production and stocking decisions. These results provide managers with actionable insights into which variables drive effective supplementation and re-evaluate their management strategies. These results highlight how the framework can capture both the handling of complexity and interpretability, two qualities essential for closing the research–implementation gap in conservation.

A practical consideration of the IPM–DRL framework is that, while the DRL model is trained under the demographic process uncertainties characterized through posterior parameter distributions, it is trained on the true latent population sizes rather than their estimates. This design choice reflects a focus on decision optimization under demographic and environmental process uncertainty and relies on the inferential model for reducing the state uncertainty. Future work could extend this framework by introducing an explicit heterogeneous observation process in the simulation and providing the DRL model with summary indices of those observations, enabling it to learn models with both process and state uncertainty. Our general approach could also be modified to incorporate improvements on the population genetics model since our assumptions may not fully capture what is occurring in the wild. For example, the genetics model could be modified to better reflect RGSM biology (e.g., high variance in reproductive success) (42,43) and the accumulative effects of inbreeding and genetic drift (SI Appendix A). These assumptions may inflate effective size estimates and influence comparisons among management strategies. Despite these limitations, the decision framework is broadly applicable to a diversity of systems and enables balancing tradeoffs among complex ecological and evolutionary processes.

Although our implementation accounted for uncertainty in the demographic parameters during DRL training, it did not explicitly model learning about those parameters through time. In classical active adaptive management models, decision-makers can reduce uncertainty about ecological processes by observing system responses to actions and adjusting models and future actions accordingly (10). While DRL agents can incorporate active learning via recurrent architectures or frame stacking (44), our value of information analysis suggests this is unnecessary here. Resolving uncertainty in winter mortality and larval carrying capacity yielded minimal performance gains, indicating that a passive adaptive strategy, periodically updating parameter distributions with new monitoring data, is sufficient. This aligns with findings in other systems that the benefits of active adaptive management often do not justify the added complexity of incorporating active learning (45).

As conservation management grows increasingly data-rich, the challenge lies in converting information into actionable, adaptive decisions. Our framework can bridge this gap by turning complex ecological data into transparent and defensible guidance. As algorithms for solving high-dimensional dynamic problems advance, our framework provides a scalable pathway to increase managers’ adoption of systematic conservation planning for endangered species.

## Method

### Simulation model

Our discrete population model describes how RGSM abundance and genetic diversity changes with the amount of hatchery supplementation. The simulations alternate between spring and fall seasons where production and stocking distribution decisions are made, respectively. The demographic component models the age-structured populations in the three reaches, in which recruitment and survival rates are a function of spring flow volume and summer river drying, respectively (SI1 Appendix A). The parameter estimates of the demographic model were derived from an IPM described and fit in Yackulic et al. (2022) that integrated multiple data sources. If the population density in any reach falls below a quasi-extinction threshold identified through expert elicitation (SI1, Appendix E) at the start of the spring spawning season, we assume that spawners would not be able to find each other, leading to low recruitment and eventual local extirpation within that reach. The reach remains empty until re-populated through sufficient stocking. If all populations in all three reaches are extirpated in a time step, the population is considered extinct in the wild and the agent receives no rewards over the remainder of the simulated time horizon.

The genetic component models the effective population size assuming a panmictic population across the three reaches. The effective population size fluctuates under the dynamics of density-dependent population growth with environmental stochasticity (46) and is impacted by hatchery supplementation through Ryman-Laikre effect (47). We note, however, that the single-generation Ryman–Laikre effect evaluated here may be conservative, as effective population size may be higher under multi-generation supplementation (48), and effective population size may be more negatively affected than variance effective population size (49). The full mathematical framework of the simulation is in the Supporting Information (SI1, Appendix A).

Our reward function, *R*, reflects two key trade-offs in the supplementation problem: balancing persistence against genetic impacts. Every year, after stocking in the fall and just before the next spring spawning season, following reward is given:

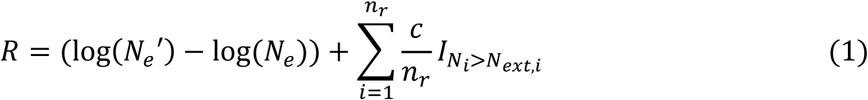

*N*′*_e_* is the effective population size after stocking, and *N_e_*is the effective population size if there had not been any stocking. *n_r_*is the number of reaches. *I_Ni_*_>*N*_*_ext_*_,*i*_ is the indicator function for whether the population size in reach *i* at the start of the spawning season, *N_i_*, exceeds its local extinction threshold, *N_ext_*_,*i*_. The relative value of persistence of RGSM, *c*, reflects the manager’s preference for prioritizing persistence over reducing genetic diversity loss. The genetic impact, log(*N_e_*′) − log(*N_e_*), focuses on reducing the log effective population size difference after stocking. This is to isolate the effect of stocking on the effective population size from the natural fluctuations of the effective population size. We set *c*=10, 4, and 1 for persistence-focused, balanced, genetics-focused objectives, respectively. The relative importance of persistence is measured as the percentage change from the mean genetic impact under the current strategy to *c*.

We also developed other derivative simulations for additional analysis. For the value of information analysis, we assumed that the agent can perfectly observe the values of some of the parameters in the model (Appendix SA). For assessing the introduction of a carry-over policy, we included the ability to store specified proportion of the produced fish to next year.

### Training and Testing Deep Reinforcement Learning model

We trained the DRL decision models using the off-policy actor–critic TD3 algorithm (50) with some modifications, which supports continuous actions necessary for proportional production and stocking decisions (SI1, Appendix F). Each training session consisted of repeated fifty-year simulations, called episodes, with state–action–reward transitions stored in the agent’s memory. At regular intervals, batches of past transitions were sampled to update the decision model (SI1, Appendix F). Multiple training sessions were run with different random seeds, and the decision model was evaluated every 1,000 episodes and saved as checkpoints. Because stochasticity in both the environment and network initialization can produce divergent learning paths, we retained only the top-performing checkpoints for each value of persistence and further evaluated over 3,000 episodes to select five representative decision models for further analyses.

For each selected agent, we computed mean local persistence rates and the harmonic mean of effective population size across 3,000 episodes. For comparison, we also evaluated several heuristic strategies: no supplementation, maximum supplementation, the current strategy, and manual decision-making. The no-supplementation strategy stocked nothing; the maximum strategy always produced at capacity and stocked evenly across reaches; and the current strategy adjusted production inversely proportional to spring-flow forecasts and allocated stocking to reaches with mean catch-per-unit-effort below 1 (SI1, Appendix A).

## Acknowledgements

This work was supported by the Bureau of Reclamation (grant number: R24AP00307). Any use of trade, firm, or product names is for descriptive purposes only and does not imply endorsement by the US Government. We thank the many individuals who contributed to the long-term datasets included in this analysis. We further thank Megan Osborne, Thomas Turner, and Thomas Archdeacon, Michael Ford, and Jaime Ashander for their helpful comments on earlier versions of this manuscript, as well as members of the expert elicitation panel.

## Author contributions

H.Y., C.Y., A.L., C.W., and K.P., designed research, performed research, analyzed data, and wrote the paper.

The authors declare no conflict of interest

## References

1. CBD. Monitoring framework for the Kunming-Montreal Global Biodiversity Framework. Conference Of the Parties to the Convention on Biological Diversity Fifteenth meeting [Internet]. 2022. Available from: https://www.cbd.int/doc/c/e6d3/cd1d/daf663719a03902a9b116c34/cop-15-l-25-en.pdf

2. Game ET, Meijaard E, Sheil D, Mcdonald-Madden E. Conservation in a wicked complex world; challenges and solutions. Conserv Lett. 2014;7(3):271–7. doi:10.1111/conl.12050

3. Iacona GD, Avery-Gomm S, Maloney RF, Brazill-Boast J, Crouse DT, Drew CA, et al. Hurdles to developing quantitative decision support for Endangered Species Act resource allocation. Frontiers in Conservation Science. 2022;3(October):1–9. doi:10.3389/fcosc.2022.1002804

4. Parker L. The War over the Delta Smelt: Balancing the Endangered Species Act with the Human Interest. University of Denver Water Law Review. 2014;18.

5. Wienhues A, Baard P, Donoso A, Oksanen M. The ethics of species extinctions. Cambridge Prisms: Extinction. 2023;1. doi:10.1017/ext.2023.21

6. Martin TG, Burgman MA, Fidler F, Kuhnert PM, Low-Choy S, Mcbride M, et al. Eliciting Expert Knowledge in Conservation Science. Conservation Biology. 2012;26(1):29–38. doi:10.1111/j.1523-1739.2011.01806.x PubMed PMID: 22280323.

7. Addison PFE, Rumpff L, Bau SS, Carey JM, Chee YE, Jarrad FC, et al. Practical solutions for making models indispensable in conservation decision-making. Divers Distrib. 2013;19(5–6):490–502. doi:10.1111/ddi.12054

8. Dubois NS, Gomez A, Carlson S, Russell D. Bridging the research-implementation gap requires engagement from practitioners. Conserv Sci Pract. 2019;2:e134. doi:DOI: 10.1111/csp2.134

9. Knight AT, Cowling RM, Rouget M, Balmford A, Lombard AT, Campbell BM. Knowing but not doing: Selecting priority conservation areas and the research-implementation gap. Conservation Biology. 2008. p. 610–7. doi:10.1111/j.1523-1739.2008.00914.x

10. Chadès I, Pascal L V., Nicol S, Fletcher CS, Ferrer-Mestres J. A primer on partially observable Markov decision processes (POMDPs). Methods Ecol Evol. 2021;12(11):2058–72. doi:10.1111/2041-210X.13692

11. Abadi F, Barbraud C, Gimenez O. Integrated population modeling reveals the impact of climate on the survival of juvenile emperor penguins. Glob Chang Biol. 2017;23(3):1353–9. doi:10.1111/gcb.13538 PubMed PMID: 27770507.

12. Riecke T V., Williams PJ, Behnke TL, Gibson D, Leach AG, Sedinger BS, et al. Integrated population models: Model assumptions and inference. Methods Ecol Evol. 2019;10(7):1072–82. doi:10.1111/2041-210X.13195

13. Dobson ADM, Milner-Gulland EJ, Aebischer NJ, Beale CM, Brozovic R, Coals P, et al. Making Messy Data Work for Conservation. One Earth. 2020;2(5):455–65. doi:10.1016/j.oneear.2020.04.012

14. Schaub M, Kéry M. Integrated Population Models: Theory and Ecological Applications with R and JAGS. Academic Press; 2021.

15. van den Berg J, Patil S, Alterovitz R. Efficient Approximate Value Iteration for Continuous Gaussian POMDPs. AAAI. 2012;1832–8. doi:10.1609/aaai.v26i1.8371

16. Zhou E, Fu MC, Marcus SI. Solving Continuous-State POMDPs via Density Projection. IEEE Trans Automat Contr. 2010;55(5):1101–16.

17. Chadès I, McDonald-Madden E, McCarthy MA, Wintle B, Linkie M, Possingham HP. When to stop managing or surveying cryptic threatened species. Proc Natl Acad Sci U S A. 2008;105(37):13936–40. doi:10.1073/pnas.0805265105 PubMed PMID: 18779594.

18. Rout TM, Hauser CE, Possingham HP. Optimal adaptive management for the translocation of a threatened species. Ecological Applications. 2009;19(2):515–26. doi:10.1890/07-1989.1 PubMed PMID: 19323207.

19. Waring TK, Somers VLJ, McCarthy MA, Baker CM. When to monitor or control: Informed invasive species management using a partially observable Markov decision process (POMDP) framework. Methods Ecol Evol. 2024;15(9):1667–76. doi:10.1111/2041-210X.14374

20. Silver D, Hubert T, Schrittwieser J, Antonoglou I, Lai M, Guez A, et al. A general reinforcement learning algorithm that masters chess, shogi, and Go through self-play. Science (1979). 2018;362(6419):1140–4. doi:10.1126/science.aar6404 PubMed PMID: 30523106.

21. Sharma AK, Kunkel J. How to Train Your Robot with Deep Reinforcement Learning – Lessons We’ve Learned. arXiv preprint. 2021;1–22. doi:10.1177/ToBeAssigned

22. Silvestro D, Goria S, Sterner T, Antonelli A. Improving biodiversity protection through artificial intelligence. Nat Sustain. 2022;5(5):415–24. doi:10.1038/s41893-022-00851-6

23. Equihua J, Beckmann M, Seppelt R. Connectivity conservation planning through deep reinforcement learning. Methods Ecol Evol. 2024;15(4):779–90. doi:10.1111/2041-210X.14300

24. Bestgen KR, Platania SP. Status and Conservation of the Rio Grande Silvery Minnow, Hybognathus amarus. Southwest Nat. 1991;36(2):225–32.

25. Archdeacon TP, Diver-Franssen TA, Bertrand NG, Grant JD. Drought results in recruitment failure of Rio Grande silvery minnow (Hybognathus amarus), an imperiled, pelagic broadcast-spawning minnow. Environ Biol Fishes. 2020;103(9):1033–44. doi:10.1007/s10641-020-01003-5

26. Archdeacon TP, Diver TA, Reale JK. Fish Rescue during Streamflow Intermittency May Not Be Effective for Conservation of Rio Grande Silvery Minnow. Water (Switzerland). 2020;12(12). doi:10.3390/w12123371

27. Kelly S, Augusten I, Mann J, Katz L. History of the Rio Grande Reservoirs in New Mexico: Legislation and litigation. Nat Resour J. 2007;47(3):525–613.

28. Archdeacon TP, Dudley RK, Remshardt WJ, Knight W, Ulibarri M, Gonzales EJ. Hatchery supplementation increases potential spawning stock of Rio Grande Silvery Minnow after population bottlenecks. Trans Am Fish Soc. 2023;152(2):187–200. doi:10.1002/tafs.10398

29. Osborne MJ, Dowling TE, Scribner KT, Turner TF. Wild at heart: Programs to diminish negative ecological and evolutionary effects of conservation hatcheries. Biol Conserv. 2020;251(August):108768. doi:10.1016/j.biocon.2020.108768

30. Yackulic CB, Archdeacon TP, Valdez RA, Hobbs M, Porter MD, Lusk J, et al. Quantifying flow and nonflow management impacts on an endangered fish by integrating data, research, and expert opinion. Ecosphere. 2022;13(9):1–22. doi:10.1002/ecs2.4240

31. Osborne MJ, Archdeacon TP, Yackulic CB, Dudley RK, Caeiro-Dias G, Turner TF. Genetic erosion in an endangered desert fish during a megadrought despite long-term breeding. Conservation Biology. 2024;38(1):e14154.

32. Bennett JR, Maxwell SL, Martin AE, Chadès I, Fahrig L, Gilbert B. When to monitor and when to act: Value of information theory for multiple management units and limited budgets. Journal of Applied Ecology. 2018;55(5):2102–13. doi:10.1111/1365-2664.13132

33. Williams B, Brown E. Adaptive Management: The U.S. Department of the Interior Applications Guide. 2012. 1–17 p.

34. Walters C. Optimum Escapements in the Face of Alternative Recruitment Hypothesis. Canadian Journal of Fisheries and Aquatic Sciences. 1981;38:678–89.

35. McMillan JR, Morrison B, Chambers N, Ruggerone G, Bernatchez L, Stanford J, et al. A global synthesis of peer-reviewed research on the effects of hatchery salmonids on wild salmonids. Fish Manag Ecol. 2023 Oct 1;30(5):446–63. doi:10.1111/FME.12643;ISSUE:ISSUE:DOI

36. Holden MH, Akinlotan MD, Binley AD, Cho FHT, Helmstedt KJ, Chadès I. Why shouldn’t I collect more data? Reconciling disagreements between intuition and value of information analyses. Methods Ecol Evol. 2024 Sep 1;15(9):1580–92. doi:10.1111/2041-210X.14391

37. Hemming V, Camaclang AE, Adams M, Burgman M, Carbeck K, Carwardine J, et al. An introduction to decision science for conservation. Conservation Biology. 2021;(January 2021):1–16. doi:10.1111/cobi.13868

38. Gibson FL, Rogers AA, Smith ADM, Roberts A, Possingham H, McCarthy M, et al. Factors influencing the use of decision support tools in the development and design of conservation policy. Environ Sci Policy. 2017;70:1–8. doi:10.1016/j.envsci.2017.01.002

39. Witzenberger KA, Hochkirch A. Ex situ conservation genetics: a review of molecular studies on the genetic consequences of captive breeding programmes for endangered animal species. Biodiversity and Conservation 2011 20:9. 2011 May 26;20(9):1843–61. doi:10.1007/S10531-011-0074-4

40. Ruiz DM, Tinker MT, Tershy BR, Zilliacus KM, Croll DA. Using meta-population models to guide conservation action. Glob Ecol Conserv. 2021 Aug 1;28:e01644. doi:10.1016/J.GECCO.2021.E01644

41. Knight AT, Cowling RM, Rouget M, Balmford A, Lombard AT, Campbell BM. Knowing but not doing: Selecting priority conservation areas and the research-implementation gap. Conservation Biology. 2008;22(3):610–7. doi:10.1111/j.1523-1739.2008.00914.x PubMed PMID: 18477033.

42. Osborne MJ, Caeiro-Dias G, Turner TF. Man-Made Barriers and Augmentation Drive Spatial and Temporal Trends of Genetic Diversity and Effective Population Size in a Riverine Fish. Mol Ecol. 2026 May 1;35(10):e70380. doi:10.1111/MEC.70380;WGROUP:STRING:PUBLICATION

43. Turner TF, Osborne MJ, Moyer GR, Benavides MA, Alò D. Life history and environmental variation interact to determine effective population to census size ratio [Internet]. doi:10.1098/rspb.2006.3677

44. Hausknecht M, Stone P. Deep recurrent q-learning for partially observable MDPs. AAAI Fall Symposium – Technical Report. 2015;FS-15-06:29–37.

45. Springborn MR. Risk aversion and adaptive management: Insights from a multi-armed bandit model of invasive species risk. J Environ Econ Manage. 2014;68(2):226–42. doi:10.1016/j.jeem.2014.05.004

46. Myhre AM, Engen S, Sæther BE. Effective size of density-dependent populations in fluctuating environments. Evolution (N Y). 2016;70(11):2431–46. doi:10.1111/evo.13063 PubMed PMID: 27624411.

47. Ryman N, Laikre L. Effects of Supportive Breeding on the Genetically Effective Population Size. Conservation Biology. 1991;5(3):325–9. doi:10.1111/j.1523-1739.1991.tb00144.x

48. Wang J, Ryman N. Genetic effects of multiple generations of supportive breeding. Conservation Biology. 2001;15(6):1619–31. doi:10.1046/j.1523-1739.2001.00173.x

49. Ryman N, Jorde PE, Laikre L. Supportive Breeding and Variance Effective Population Size. Conservation Biology. 1995;9(6):1619–28. doi:10.1046/j.1523-1739.1995.09061619.x

50. Fujimoto S, Van Hoof H, Meger D. Addressing Function Approximation Error in Actor-Critic Methods. 35th International Conference on Machine Learning, ICML 2018. 2018;4:2587–601.

