## Supplemental Information 1 for "Bridging Ecological Inference and Decision Optimization for Conservation Using Artificial Intelligence"

**Supporting Information 1**

**Content**

**Appendix A. Simulation model.**

**A1. Simulation description**

**A2. Initialization**

**A3. Current strategy**

**A4. Derivative simulations**

**Appendix B. Integrated population modeling**

**Appendix C. Modeling hydrological process and forecast**

**Appendix D. Larval carrying capacity index prediction**

**Appendix E. Elicitation of local quasi-extinction threshold**

**Appendix F. Deep reinforcement learning**

**Reference**

**Appendix A. Simulation model.**

**A1. Simulation description**

The simulation model for the supplementation of Rio Grande silvery minnow (RGSM) is a Partially Observable Markov Decision Process (POMDP) model, where the true state of the environment is only imperfectly observed by the conservation manager. The simulation describes the population dynamics of RGSM on a seasonal basis (spring to fall and fall to spring) in terms of survival and reproduction rates that depend on hydrological conditions. The population is structured by age (0 and 1+ year fish) and by space (Angostura, Isleta, and San Acacia reach). The demographic component of the simulation largely follows the IPM model in Yackulic et al. (2022).

The sub-population in a given reach is assumed to be locally extinct if the density falls below a pseudo-extinction threshold at the start of the spawning season. A sub-population can be restored if the agent, or the virtual manager, adds enough fish in a future year to exceed the pseudo-extinction threshold. If all three sub-populations go extinct at once, the population is considered fully extinct in the wild, and the simulation terminates.

The simulation model also keeps track of the dynamics of the effective population size in the wild population. The wild effective population size depends on the density-dependent fluctuations in the population size, as well as the total amount of fish stocked through supplementation. Although dispersal among reaches is assumed to be negligible, the sub-populations remain genetically indistinguishable (Osborne et al., 2020).

More than two decades of genetic monitoring indicate that genetic diversity in the RGSM population has remained well maintained through time despite repeated demographic contractions (Osborne et al. 2020). This outcome reflects deliberate genetic planning embedded in the supplementation program, including the routine use of wild-origin broodstock and periodic broodstock refreshment across the spatial extent of the wild population to represent standing genetic variation. Hatchery production is frequently based on collections of wild fish, often as eggs that are hatched by the thousands in captivity and stocked later, and in some years also incorporates salvaged wild individuals brought into the hatchery during emergency rescue operations. By design, these practices maintain strong genetic continuity between hatchery and wild components, limiting divergence and minimizing the effects of hatchery selection and genetic drift associated with small broodstock. Consequently, supplementation can function as a genetic and demographic buffer during severe bottleneck events, when effective population size and rare alleles could otherwise be lost rapidly due to catastrophic mortality and high variance in reproductive success.

Each discrete step of the simulation alternates between spring and fall seasons. The conservation manager takes actions on how many fish to produce in the spring and how to distribute the produced fish across the three reaches in the fall. The goal of hatchery supplementation is to maximize the persistence probability of the wild population and the genetic impact to the effective population size. Figure S1 provides a conceptual overview of the annual decision cycle, illustrating the timing of production and stocking decisions, the monitoring data available to managers at each stage, and key environmental and demographic processes influencing population dynamics between decisions modeled in the simulation.

Formally, the POMDP model is described as a 7-tuple

$$< S, A, T, R, \Omega, O, \gamma>$$

where

- $S$ is a set of states,
- $A$ is a set of actions,
- $T$ is a transition model between states,
- $R : S \times A \to R$ is the reward function,
- $\Omega$ is a set of observations,
- $O$ is an observation model, and
- $\gamma$ ∈ [0, 1) is the discount factor (fixed to 0.99).

We described the simulation model by defining each element of the POMDP tuple in the context of our supplementation model.

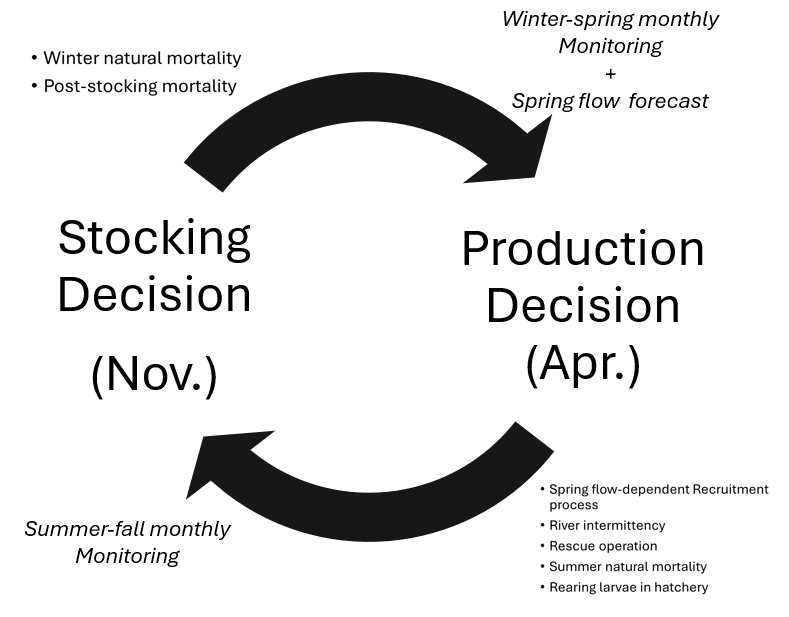

Figure S1. Conceptual diagram of the annual supplementation decision cycle for Rio Grande silvery minnow. Production decisions are made in spring (April) with the availability of winter–spring monitoring data together with spring flow forecasts, whereas stocking decisions are made in fall (November) with the availability of summer–fall monitoring. Arrows indicate the seasonal sequence of management actions and key ecological processes, including flow-dependent recruitment, river intermittency, rescue operations, natural mortality, and post-stocking mortality, that influence population dynamics between decision points.

**A1.1 States (**$\boldsymbol{S)}$

State variables describe the demographic, genetic, and hydrological components of the environment. Subscript r ∈ {1, 2, 3} describes different reaches (1=Angostura, 2=Isleta, 3=San Acacia).

- $N_{r,0}$: Population size of age 0 fish in reach $r$
- $N_{r,0CF}$: Population size of wild age 0 fish in reach $r$. This is the number of wild age 0s – in years with stocking this can also be thought of the counterfactual to the number of age0s after stocking in the fall.
- $N_{r,1}$: Population size of age 1+ fish in reach $r$
- $N_{h}$: Number of fish in the hatchery
- $q$: Amount of total spring flow at Otowi stream gage from March 1st to July 31st.
- $N_{e}$: Effective population size of the wild population
- *t*: Binary season indicator (0 = spring, 1 = fall).
- $\theta$: demographic parameter values.

**A1.2 Observations (**$\boldsymbol{\Omega}$**)**

Observation variables describe the information available to the manager about the current state of the environment. We assumed that $N_{r,0}$, $N_{r,0CF}$, $N_{r,1}$, $N_{h}$, and $N_{e}$ are directly observable at every timestep. Conversely, we assumed that the demographic process parameters, $\theta$, are not observable. In the spring, the manager only observes the forecast of the spring flow, which is the actual spring flow value plus the forecast bias. We denoted the spring flow forecast observation variable as $\hat{q}$. In the fall season, the spring flow in that year becomes directly observable.

**A1.3 Action (**$\boldsymbol{A}$**)**

Action variables describe the management decisions made each year, including how many fish to produce in the spring and how to allocate the hatchery-reared fish across the three river reaches in the fall. An action $a\in A$ is a vector of length 4,

$$a =(p_{p}, p_{a}, p_{i}, p_{s})$$

Where $p_{p}$is the proportion of fish produced in spring ($0\leq p_{p}\leq1$), $p_{a}, p_{i},$ and $p_{s}$ are the proportions of produced fish stocked into Angostura, Isleta, and San Acacia reaches, respectively ($p_{a}+p_{i}+p_{s}=1$). Note that only the production action matters in determining the state transition from spring to fall, and only the stocking distribution from fall to spring. Masking unnecessary action variables in a given season during the training process of the DRL model is outlined in Appendix E.

**A1.4 State transition (**$\boldsymbol{T}$**)**

The state transition model $\boldsymbol{T}$ describes how the state variables transition between the steps. The transition rules for many variables depend on the season. Note that the “+” superscript indicates the state variable in the next step.

**A1.4.1 Season (t)**

The season indicator variable, *t*, is a binary variable alternating between 0 and 1 each step, characterizing the change between spring ($t = 0$) and fall seasons ($t = 1$).

$$\begin{aligned} t^{+}=1-t \#\left( 1 \right) \end{aligned}$$

The exact date of the year for the two points is March 31^st^ and November 1^st^ for spring and fall, respectively.

**A1.4.2 Wild population size**

a) Spring ($t=0$)

The transition of population size from spring to fall is determined by the recruitment process from April through June, and summer mortality, which is affected by river drying events and rescue actions, where managers relocate the fish trapped in the pools on dried riverbeds.

$$\begin{aligned} N_{r,0}^{+}=P_{r}e^{-124M_{r0}}\left( \left( 1-\delta_{r} \right)+\tau\left( \Delta_{\delta_{r}} \right)+\left( 1-\tau\right)r_{0}\Delta_{\phi} \right) \# \end{aligned}\left( 2 \right)$$

$$\begin{aligned} N_{r,1}^{+}=\left( N_{r,0}+N_{r,1} \right)e^{-215M_{r1}}\left( \left( 1-\delta_{r} \right)+\tau\delta_{r}+\left( 1-\tau\right)r_{1}\phi\right) \#\left( 3 \right) \end{aligned}$$

$P_{r}$ is the recruitment size in reach $r, \tau$ is the proportion of fish that successfully moves out of the dried parts of the river, and $M_{r0}\mathrm{and} M_{r1}$are the daily spring-summer mortality rate for age 0 and age 1+ fish, respectively. $\delta_{r}$ is the proportion of the river that dried at least once by November 1st that year and $\Delta_{\delta_{r}}$ is the proportion of the river that dried at least once between July 1st and November 1st.$\phi$ is the weighted average survival rate of rescued RGSM at any point through November 1st, and $\Delta_{\phi}$ is the weighted average survival rate of rescued RGSM past July 1st up to November 1st. The value of 124 in Equation (2) corresponds to the number of days between July 1st, when larvae are recruited to age 0, and November 1st, when the next timestep begins ($t=1$). The value of 215 in Equation (3) is the number of days between March 31st ($t=0$) to November 1st ($t=1$). See Yackulic et al. (2022) for more information on these equations.

The river drying-related parameters, $\delta_{r}$ and $\Delta_{\delta_{r}}$ are random variables with beta distributions fitted to past river drying data for Isleta and San Acacia reaches ($r=2,3$) (Appendix C).

$$\begin{aligned} \delta_{r}\sim Beta\left( \alpha_{\delta},\beta_{\delta} \right)\#\left( 4 \right) \end{aligned}$$

$$\begin{aligned} \Delta_{\delta_{r}}\sim Beta\left( \alpha_{\Delta_{r}},\beta_{\Delta_{r}} \right)\#\left( 5 \right) \end{aligned}$$

The estimates for $\alpha_{\delta},\beta_{\delta}$, $\alpha_{\Delta_{r}},{\mathrm{and} \beta}_{\Delta_{r}}$ are, 0.87, 4.51, 1.08, and 0.57 for Isleta, and 1.60, 2.14, 1, and 0.60 for San Acacia, respectively (Appendix C). Angostura reach is assumed not to dry, and those parameters are set to 0. This assumption was true during the period studied by Yackulic et al. (2022) during which all demographic parameters were estimated, however we note that drying has occurred at times prior to and after their study period and future work could relax this assumption.

Daily mortality rates, $M_{r0}$ and $M_{r1}$, are further characterized by parameter $m_{0}\mathrm{and}m_{1}$as follows:

$$\begin{aligned} M_{r0}=e^{m_{0}}\quad\#\left( 6 \right) \end{aligned}$$

$$\begin{aligned} M_{r1}=e^{m_{1}}\quad\#\left( 7 \right) \end{aligned}$$

The recruitment size, $P_{r}$, is defined with Beverton-Holt equation.

$$\begin{aligned} P_{r}=\frac{\alpha S_{r}}{1+\alpha S_{r}/\kappa_{r}}\#\left( 8 \right) \end{aligned}$$

where $\alpha$ is the mean number of age 0 RGSM produced by each effective spawner, $S_{r}$ is the number of effective spawners, $\kappa_{r}$ is the larval carrying capacity.

The number of effective spawner in reach $r$, $S_{r}$, is defined as

$$\begin{aligned} S_{r}=N_{r0}+\beta_{2}N_{r1}\#\left( 9 \right) \end{aligned}$$

where $\beta_{2}$ is the relative spawning capability of age 2+ fish in relation to age 1 fish. Note that $N_{r0}$ would denote the population of age 1 and $N_{r1}$ the population of age 2+ at the moment of spawning.

The larval carrying capacity at reach $r,\kappa_{r}$, is defined as

$$\begin{aligned} \kappa_{r}=e^{\beta\left( L_{r}-\bar{L}_{r} \right)+\epsilon_{r}}\#\left( 10 \right) \end{aligned}$$

where $L_{r}$ is the larval carrying capacity index, $\bar{L}_{r}$ is the mean of past estimates of larval carrying capacity indexes (=0.652), and $\epsilon_{r}$ is the stochastic reach-specific effect defined by the normal distribution.

$$\begin{aligned} \epsilon_{r}\sim\mathcal{N}\left( \mu_{Lr},\sigma_{L} \right)\#\left( 11 \right) \end{aligned}$$

The larval carrying capacity index, $L_{r}$, is a function of the spring flow of each reach that is defined by a general additive model fit to the hydrograph and monitoring data (Appendix D).

The counterfactual age 0 population is the same as the regular age 0 population at the start of fall.

$$\begin{aligned} N_{r0CF}^{+}=N_{r0}^{+}\#\left( 12 \right) \end{aligned}$$

b) Fall ($t=1$)

In the fall season, the population size transition involves the stocking of age 0 fish produced from the hatchery that spring, survival of the population through the winter, and the local extinction process. If the total population size in a reach by the start of the spring season is below the quasi-extinction threshold for that reach ($N_{ext,r}$), we assumed that the spawner density in a reach is too small to successfully spawn and the reach’s population goes to extinction. The quasi-extinction threshold for a reach is defined as the length of that reach multiplied by the density quasi-extinction density threshold $d_{ext}$, which was elicited by a panel of experts (Appendix E).

$$\begin{aligned} N_{r,0}^{+}=\left\{ \begin{aligned} \left( N_{r,0}+N_{h}p_{r}\phi_{w} \right)e^{-150M_{w,r}} \mathrm{if}\left( N_{r0}+N_{h}p_{r}\phi_{w}+N_{r1} \right)e^{-150M_{w}}>N_{ext,r}) \\ 0 otherwise \end{aligned} \right.\#\left( 13 \right) \end{aligned}$$

$$\begin{aligned} N_{r,1}^{+}=\left\{ \begin{aligned} N_{r,1}e^{-150M_{w,r}} \mathrm{if}\left( N_{r0}+N_{h}p_{r}\phi_{w}+N_{r1} \right)e^{-150M_{w}}>N_{ext,r}) \\ 0 otherwise \end{aligned} \right.\#\left( 14 \right) \end{aligned}$$

$p_{r}$ is the proportion of the fish produced in the spring that is stocked in reach $r$, and $\phi_{w}$ is the proportion of stocked fish that survive the stocking event. $M_{w}$ is the daily mortality rate in the winter months from November 2^nd^ through March 31^st^, characterized by $m_{w}$.

$$\begin{aligned} M_{w}=e^{m_{w,r}}\#\left( 15 \right) \end{aligned}$$

The value of 150 is the number of days in between those two dates. The transition of counterfactual age 0 population differs from the age 0 population in the fall, as it would not account for the stocking.

$$\begin{aligned} N_{r,0CF}^{+}=\left\{ \begin{aligned} N_{r,0}e^{-150M_{w}} if \left( N_{r0}+N_{r1} \right)e^{-150M_{w}}>N_{ext,r}) \\ 0 otherwise \end{aligned} \right.\#\left( 16 \right) \end{aligned}$$

Thus, Equation (16) denotes the age 1 population at the start of the spring season that was born in the wild.

**A1.4.3 Spring flow volume (**$\boldsymbol{q}$**)**

$q$ is the total amount of flow in spring at Otowi from March 1st to July 31st and it mainly serves as determining the larval carrying capacity in the demographic process. Otowi reach is located upstream of Angostura reach, where the spring flow forecast from the Natural Resources Conservation Service (NRCS) is made. The spring flow is drawn each year from a normal distribution that was fitted to the historical hydrological data from 1997 to 2024 (Appendix C).

$$\begin{aligned} q^{+}\left\{ \begin{aligned} \mathrm{logistic}\left( \mathcal{N}\left( \mu_{q},\sigma_{q} \right) \right){(q_{max}-q}_{min})+q_{min} if t=1 \\ q if t=0 \end{aligned} \right.\#\left( 17 \right) \end{aligned}$$

${q_{max} \mathrm{and}q}_{min}$ are 110% of the maximum and 90% of the minimum spring flow observed at Otowi between 1997 and 2024, respectively. The logistic transformed $\mu_{q} \mathrm{and}\sigma_{q}$ estimates are 957.2 and 468 thousand acre-ft, respectively (Appendix C). Spring flow at the three main reaches where RGSM are found are derived from $q$ variable by subtracting a reach-specific constant, which is then used to calculate the reach-specific larval carrying capacity index (Appendix C).

**A1.4.4 Hatchery population size (**$\boldsymbol{N}_{\boldsymbol{h}}$**)**

$N_{h}$ is the number of fish in the hatchery for stocking. The hatchery population increases from production in the spring ($t=0$). The hatchery population is emptied to 0 at $t=0$

$$\begin{aligned} {N_{h}}^{+}\left\{ \begin{aligned} 0 if t=1 \\ mp_{p} \mathrm{if} t=0 \end{aligned} \right.\#\left( 18 \right) \end{aligned}$$

$m$ is the maximum capacity of fish that the hatchery can hold (=200,000) and $p_{p}$ is the production action.

**A1.4.5 Wild effective population size (**$\boldsymbol{N}_{\boldsymbol{e}}$**)**

Effective population size measures the rate of genetic diversity loss in a population. The wild effective population size depends on two factors in our model. First, it is affected by the demographic size of the population. To model this, we used the equation that describes the effective population size of a fluctuating density-dependent population under environmental stochasticity from Myhre et al., (2016). Second, it is affected by the hatchery supplementation through the admixture process. We followed Ryman & Laikre, (1991) that describes the wild effective population size after stocking (Eq. 30).

In the effective population size state variable, we only measured the effective population sizes defined by the fluctuations in the wild population size. The effect of stocking in the fall on the effective population size is modeled in the reward system, and the effective population size is only updated in the spring post-spawning.

In spring, the effective population size of the wild population, $N_{e}$, at the end of spring transitions is

$$\begin{aligned} N_{e}^{+}=\frac{N_{r0CF}}{\int\frac{\sigma_{dg}^{2}\left( N_{r0CF},N_{r1} \right)}{\lambda^{2}\left( N_{r0CF},N_{r1} \right)}f\left( \boldsymbol{u} \right)d\boldsymbol{u}}\#\left( 19 \right). \end{aligned}$$

$\boldsymbol{u}$ is the vector of environmental variables, namely, the larval carrying capacity ($\kappa_{r}$) and proportions of river drying in the summer ($\delta_{r}$), and $f\left( \boldsymbol{u} \right)$is the probability density of $\boldsymbol{u}$. The integral was calculated numerically by discretizing the environmental variables. $\lambda$ and $\sigma_{dg}^{2}$ denote the expected population growth rate and the demographic variance of population growth rate, respectively. The two variables are defined as follows from Myhre et al. (2016):

$$\begin{aligned} \lambda=s_{a}+b_{w}/2\#\left( 20 \right) \end{aligned}$$

$$\begin{aligned} \sigma_{dg}^{2}=s_{a}\left( 1-s_{a} \right)+b_{w}/4+\sigma_{b_{w}}^{2}/4\#\left( 21 \right) \end{aligned}$$

$s_{a}$ is the expected survival rate of adults (age 1+), $b_{w}$ is the expected number of recruits (offspring that make it to age 1) produced per spawner, and $\sigma_{b_{w}}$ is the variance in the number of recruits produced per spawner. The adult survival rate $s_{a}$ is further defined as the geometric mean of adult reach-specific survival rates, $s_{a,r}$, across the three reaches.

$$\begin{aligned} s_{a,r}=e^{-215\mu_{m_{r,0}}-150\mu_{m_{r,w}}}\left( \left( 1-\delta_{r} \right)+\tau\delta_{r}+\left( 1-\tau\right)r_{1}\phi_{r} \right)\#\left( 22 \right) \end{aligned}$$

Expected number of recruits per spawner is calculated by spawner age class and averaged in a weighted manner.

$$\begin{aligned} b_{w,0}=\sum_{r} \left( \frac{\alpha\sum_{r} N_{r,0}}{1+\alpha S_{r}/\kappa_{r}}e^{-124m_{r,0}-150\mu_{m_{r,w}}}\left( \left( 1-\delta_{r} \right)+\tau\left( \delta_{r}\Delta_{\delta_{r}} \right)+\left( 1-\tau\right)r_{0}\left( \Delta_{\phi_{r}} \right) \right) \right)/\sum_{r} N_{r,0}\#\left( 23 \right) \end{aligned}$$

$$\begin{aligned} b_{w,1}=\sum_{r} \left( \frac{\alpha\beta_{2}\sum_{r} N_{r,1}}{1+\alpha S_{r}/\kappa_{r}}e^{-124m_{r,1}-150\mu_{m_{r,w}}}\left( \left( 1-\delta_{r} \right)+\tau\left( \delta_{r}\Delta_{\delta_{r}} \right)+\left( 1-\tau\right)r_{0}\left( \Delta_{\phi_{r}} \right) \right) \right)/\sum_{r} N_{r,1}\#\left( 24 \right) \end{aligned}$$

$$\begin{aligned} b_{w}=\frac{N_{0}b_{w,0}+N_{1}b_{w,1}}{N_{0}+N_{1}}\#\left( 25 \right) \end{aligned}$$

Lastly, the variance in the number of recruits per spawner, $\sigma_{b_{w}}$, is derived by calculating the expected value of the squared difference between $b_{w}$ and $B_{w}$, the number of recruits per spawner given the number of age 0 surviving to July 1^st^ per spawner, $H$ ($E(H)=\alpha$).

$$\begin{aligned} \sigma_{b_{w}}^{2}=E\left[ \left( B_{w}\left( H \right)-b_{w} \right)^{2} \right]=\int\left( B_{w}\left( h \right)-b_{w} \right)^{2}f\left( h \right)dh\#\left( 26 \right) \end{aligned}$$

The distribution of $H$ assumed a normal distribution with the variance characterized by the mean and variance estimates of fecundity (eggs released per female; mean=3,017, variance=478) and fertilization rate (mean=0.645, variance=0.25) from Caldwell et al. (2019), assuming equal sex ratio (Osborne 2022) and the survival rate to July 1^st^ as 0.75 (Yackulic et al., 2022). The estimated variance of $H$ came out to be 94,513. Sensitivity analysis of varying this value by up to 5-fold only led to less than 2 to 3 percent changes in effective population size calculations. The integral is calculated numerically through discretizing $H$.

A limitation of our $N_{e}$ formulation is that the Myhre et al. (2016) framework assumes well-mixed reproduction and does not represent spatial structure in variance in reproductive success. The panmixia assumption itself is empirically supported, where multi-year genetic monitoring shows no significant differentiation among reaches (Osborne & Turner, 2021). However, the combination of pelagic-egg life history and diversion dams generates a sweepstakes-mismatch process in which a small fraction of spawners contributes most recruits in a given year, and spawners positioned near a diversion contribute disproportionately little because their eggs drift past the dam before recruiting (Alò & Turner, 2005; Osborne et al., 2005). This mechanism is thought to drive the very low effective-to-census-size ratios observed empirically in RGSM (on the order of 10⁻³). Our model therefore likely overestimates the effective population size in absolute terms, though relative comparisons among strategies, based on log-ratios of post- to pre-stocking effective size, should be more robust than the absolute values.

**A1.4.6 Demographic parameters (**$\boldsymbol{\theta}$**)**

The parameters that are estimated by the IPM and have defined posterior distributions are effectively state variables because they are sampled at every new iteration of the simulation. Once sampled at the start of an iteration, these parameters stay fixed until the iteration is over. The DRL model never observes these parameter values. Unlike the retrospective IPM of Yackulic et al. (2022), our forward-looking simulation does not assign fixed year-specific parameter values to future years. For parameters with year-specific estimates in the IPM, such as seasonal mortality rates, we summarized temporal variation using the cross-year means and standard deviations of their posterior estimates, and represented annual variation in the simulation through stochastic draws from these summaries. This approach preserves the magnitude of historical interannual variation while avoiding conditioning future years on specific observed years.

Table S1. Demographic parameters estimated by the IPM.

| Notation | Description | Mean | SD |
| --- | --- | --- | --- |
| $\alpha$ | mean number of offspring per spawner that survive to July 1st | 737 | 123 |
| $\beta$ | slope of the larval carrying capacity index | 9.224 | 0.964 |
| $\mu_{L,1}$ | reach-specific intercept for larval carrying capacity for Angostura | 9.751 | 0.572 |
| $\mu_{L,2}$ | reach-specific intercept for larval carrying capacity for Isleta | 12.804 | 0.532 |
| $\mu_{L,3}$ | reach-specific intercept for larval carrying capacity for San Acacia | 12.556 | 0.507 |
| $\sigma_{L}$ | random effect for larval carrying capacity | 1.528 | 0.220 |
| $\beta_{2}$ | relative reproductive contribution of age 2+ spawner | 2.001 | 0.098 |
| $\tau$ | proportion of fish that move out of a drying river segment | 0.058 | 0.019 |
| $r_{0}$ | probability age 0 fish survives in a drying river segment until rescued | 0.031 | 0.009 |
| $r_{1}$ | probability age 1+ fish survives in a drying river segment until rescued | 0.324 | 0.105 |
| $m_{0,1}$ | log summer natural daily mortality rates for Angostura for age 0 | -4.731 | 0.459 |
| $m_{0,2}$ | log summer natural daily mortality rates for Isleta for age 0 | -4.068 | 0.189 |
| $m_{0,3}$ | log summer natural daily mortality rates for San Acacia for age 0 | -5.133 | 0.441 |
| $m_{1,1}$ | summer natural daily mortality rates for Angostura for age 1+ | -4.801 | 0.383 |
| $m_{1,2}$ | summer natural daily mortality rates for Isleta for age 1+ | -4.769 | 0.226 |
| $m_{1,3}$ | summer natural daily mortality rates for San Acacia for age 1+ | -4.095 | 0.156 |
| $m_{w,1}$ | log winter natural daily mortality rates for Angostura | -5.051 | 0.405 |
| $m_{w,2}$ | log winter natural daily mortality rates for Isleta | -5.384 | 0.359 |
| $m_{w,3}$ | log winter natural daily mortality rates for San Acacia | -6.807 | 0.410 |
| $\phi_{w}$ | proportion of stocked fish that survive the stocking event | 0.154 | 0.042 |

**A1.5 Observation model (**$\boldsymbol{O}$**)**

All state variables are visible except for the parameter variables and the spring flow variable ($q$). Spring flow is only partially observable as forecast, $\hat{q}$.

$$\begin{aligned} \hat{q}=\left\{ \begin{aligned} q \mathrm{if} t=1 \\ q+\epsilon_{q} \mathrm{if} t=0 \end{aligned} \right. \#\left( 27 \right) \end{aligned}$$

$$\begin{aligned} \epsilon_{q}\mathcal{\sim N}\left( \mu_{q},\sigma_{q} \right)\#\left( 28 \right) \end{aligned}$$

Estimates for $\mu_{q},\sigma_{q}$ are 9.02, and 100.45 thousand acre-ft, respectively.

**A1.6 Reward (**$\boldsymbol{R}$**)**

The reward function defines the management objective for a single time step as a function of the current state and the action taken. In the context of the hatchery supplementation task, the reward function is defined to reflect the conservation objective of the supplementation, which is supporting long-term persistence of the RGSM population and minimizing impacts on the wild genetic diversity. Specifically, the reward function is formulated as

$$\begin{aligned} R=\left\{ \begin{aligned} \sum_{i=1}^{n_{r}} \frac{c}{n_{r}}I_{N_{i}^{+}>N_{ext,i}}+(log \left( N_{e}' \right)-\log(N_{e})) if t=1 \\ 0 if t=0 \end{aligned} \right. \#\left( 29 \right). \end{aligned}$$

$c$ is the relative value of persistence, $n_{r}$ is the number of reaches, $I_{N_{i}>N_{ext,i}}$ is the identity function for when the population size at reach $i$ at the start of spring spawning is greater than its local extinction threshold, $N_{ext,i}$, $N_{e}$ is the wild effective population size and $N_{e}'$ is the total effective population size after stocking. The relative value of persistence of RGSM, $c$, determines the manager’s emphasis on supporting persistence over reducing the decrease in effective population size. As shown in Equation (29), the reward is only conferred at the end of the fall step. The total effective population size after stocking, $N_{e}'$, is dependent on the wild and hatchery broodstock effective population sizes through the Ryman-Laikre effect.

$$\begin{aligned} N_{e}^{'}=\left\{ \begin{aligned} N_{e} \mathrm{if} x=0 \\ \frac{1}{\frac{x^{2}}{N_{e,h}}+\frac{\left( 1-x \right)^{2}}{N_{e}}} \mathrm{otherwise} \end{aligned} \right.\#\left( 30 \right) \end{aligned}$$

$N_{eh}$ is the effective population size of the hatchery broodstock, $N_{e}$ is the wild effective population size, and $x$ is the proportion of contribution of the hatchery broodstock to the next generation of spawners in the wild.

$$\begin{aligned} x=1-\frac{\sum_{r} N_{r0CF}^{+}}{\sum_{r} N_{r0}^{+}}\#\left( 31 \right) \end{aligned}$$

$$\begin{aligned} N_{eh}=N_{b}\rho_{N_{e}}\#\left( 32 \right) \end{aligned}$$

$$\begin{aligned} N_{b}=2N_{h}/R_{f}\#\left( 33 \right) \end{aligned}$$

$N_{b}$ is the number of broodstock used to produce the amount of stock-ready fish in the hatchery, $\rho_{N_{e}}$ is the mean Ne-to-N ratio in the broodstock (=0.875) estimated from Osborne et al. (2013), and $R_{f}$ is the mean number of fish produced per female broodstock (=645) estimated from the hatchery facility data.

**A2. Initialization**

The simulation always started from the fall season with no fish produced in the hatchery. Initialization of population sizes consists of first drawing the total population size between the sum of local extinction thresholds and the maximum population and 10 million in a uniform distribution. Next, the proportion of age 0 was sampled from a uniform distribution, and then the proportions for each reach from a Dirichlet distribution (Dirichlet distribution parameters set to (1,1,1)). The counterfactual age 0 population is initialized with the initialized age 0 population. Effective population size is set to 60 percent of the total population size. Spring flow rates are randomly drawn from Equation (17). The initial state was not entered into the training process for the DRL and had immaterial effect on it overall.

**A3. Current strategy**

The current strategy heuristics follow the annual supplementation plan outlined in Archdeacon, (2022) with some modifications. The current stocking decision depends on the average catch per unit effort (CPUE) values in each reach from the monitoring in October. If the CPUE goes below 1.0, the amount that would bring the expected CPUE back to 1 is stocked. Any excess fish after that is distributed using the manager’s discretion. However, comparing this strategy directly with DRL-derived policies would give advantage to DRL-derived policies as CPUE does not accurately reflect the population size due to observation errors. To make a fairer comparison, we translated this ruleset to stocking back up to the population size that 1 CPUE would correspond to, given the average catch probability estimated in Yackulic et al. (2022). If there are excess fish in the hatchery after applying this stocking rule, the rest are distributed according to the proportions of inverse of reach-level populations. If there are fewer fish in the hatchery than the rule would require, they are distributed by proportions of the amount required across reaches.

Production decision of the current strategy depends on the spring flow forecast. The decision uses the pred*i*ction from the generalized linear regression with quasi-binomial error fit to the past data on how many fish were required to stock and the spring flow volume at Otowi in the past.

$$\begin{aligned} \left( production target \right)=min\left( -0.0054\hat{q} + 2.32, m \right) \#\left( 34 \right) \end{aligned}$$

$\hat{q}$ here is in the unit of thousand acre-ft. The amount of production is capped by the current maximum capacity of the hatchery facility, $m$.

A4. Derivative simulations.

Two other simulations were used to train the DRL to analyze the effect of carry-over flexibility and perfect information on demographic parameters. In analyzing the carry-over flexibility, the action vector included an additional variable $p_{c}$, that indicated the proportion of fish to carry over to next year. The stocking distribution was applied to the leftover fish after the carry-over decision, making the proportion of fish stocked in, for example, Angostura, $p_{c}p_{a}$. In defining the hatchery population, $N_{h}$, we included an additional variable for fish that were carried over from last year. In the spring, the carried-over population would remain the same to the next step; in the fall, it would return to 0, as all carried-over fish must be stocked the following year. Equation (13) that describes winter survival process is modified to include both the newly produced and carried-over fish from last year.

In the perfect information analysis, some of the parameters are observable. We decided to analyze the value of perfect information for the parameters related to winter mortality and larval carrying capacity as these two components critically affect the population dynamics of the RGSM life cycle. For analyzing the expected value of perfect information on winter mortality, $m_{w,r}$ is observable. For analyzing the expected value of perfect information on reach-specific larval carrying capacity, $\mu_{L,r}$ is observable. The complete list of these parameters is listed in Table S1.

**Appendix B. Integrated population modeling**

We used the IPM model from Yackulic et al. (2022) to apply the IPM-DRL framework to the RGSM supplementation task. Briefly, the IPM synthesizes multiple heterogeneous data sources, including long-term catch-per-effort monitoring, fish rescue data collected during river drying, fall population size estimates, hydrological records, hatchery release records, laboratory studies on fecundity and larval survival, and structured expert elicitation, into a unified, age-structured population model for the RGSM. The model tracks dynamics in three hydrologically isolated reaches across four demographic classes (age 0, age 1, age 2 and above, and hatchery-supplemented), with key life-history transitions tied to biologically meaningful dates. To relate latent population size to catch data, the IPM includes a detailed observation model, where the predicted catch depends on capture probabilities, mesohabitat availability and preference, and the fraction of the river segment exposed to sampling. This model structure allows the IPM to partition recruitment, summer mortality, drying-related mortality and rescue, and overwinter survival while explicitly estimating uncertainty through hierarchical random-effects formulations.

Model parameters were estimated in a Bayesian framework implemented in Stan (version 2.32.6), with weakly informative priors for most parameters and informative priors for reproductive parameters derived from laboratory studies. Competing formulations of the larval carrying capacity index were compared using segment-specific R² metrics and out-of-sample forecasts for 2019 and 2020. The final selected model successfully reproduced observed catch patterns and provided posterior distributions for demographic parameters and latent population sizes. These posterior parameter samples are used to characterize demographic process uncertainty in our simulation environment, and the IPM-estimated population size from future monitoring will serve as inputs to the trained DRL model for deriving production and stocking distribution recommendations.

**Appendix C. Modeling hydrological process and forecast**

We simulated the annual spring flow using a normally distributed white noise distribution. The model was fit to the data on flow volume from April 1^st^ to July 31^st^ at the Otowi gage (08313000; USGS (2026)) for the Otowi gage from 1997 to 2024 (Fig. S2). This timeframe was chosen because the dynamics of the volume were stationary within this period. To ensure that we do not model spring flow values drastically outside of the observed values, we fit the model to a transformed version of the data. The transformation consisted of going through a min-max normalization using 90 percent of the minimum observed flow and 110 percent of the maximum observed flow, and then putting that through a logit transformation. We selected the white noise model over various autoregressive moving average alternatives because it yielded the lowest AIC, and the temporal models showed no significant parameter estimates. Figure S4 shows the comparison of the histograms of observed and modeled spring flows.

The spring flow at Angostura, Isleta, and San Acacia reaches was derived from that of the Otowi gage by subtracting the difference in the mean spring flow between the Otowi gage and other two gages (Albuquerque and San Acacia gages). The spring flow of Angostura reach was derived from the estimates for the Albuquerque gage; the spring flow of Isleta and San Acacia reaches were both derived from the estimates for the San Acacia gage. We chose this method to model the spring flows for the reaches because the regression result between the spring flows at Otowi and other reaches showed a slope of nearly 1, indicating almost a constant difference between them. The spring flows of the reaches were also clipped by 90 percent of the minimum and 110 percent of the maximum spring flow observed for their respective reaches.

We simulated the spring flow forecast that the manager observes in the spring by fitting the bias between the spring flow data and the NRCS 50% exceedance forecast data at Otowi gage to a normal distribution (Figure S3). There was no significant autocorrelation structure in the bias across the years.

The variables related to river-drying ($\delta$ and $\Delta_{\delta}$) were fit to the data from the River Eyes monitoring report, which provides daily observation measurements of the proportion of river channel that dried within the middle Rio Grande region (McKenna, 2019). We fit the parameters to a beta distribution for each Isleta and San Acacia reach. We assumed that Angostura reach does not dry because although the reach dries rarely, it did not occur during the period studied by Yackulic et al. (2022). Figure S5 shows the fitted beta distribution and the observed river-drying variables.

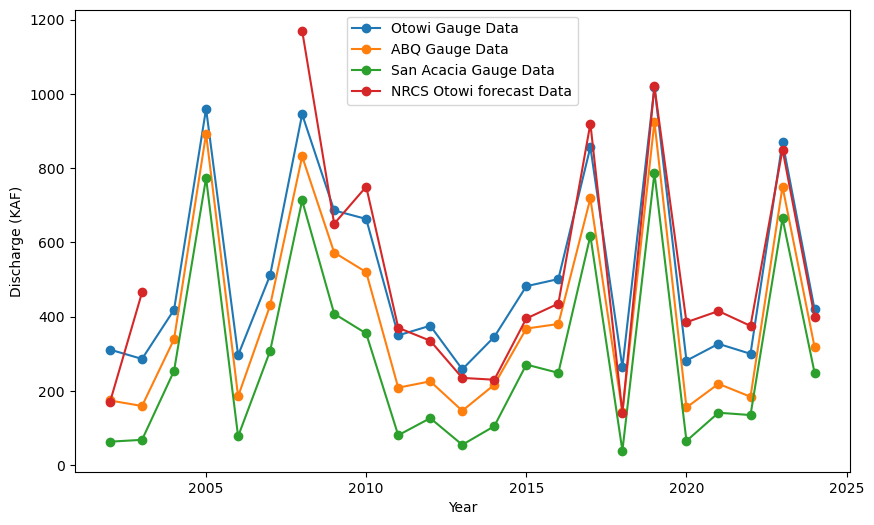

Figure S2. Annual spring flow in thousands of acre-feet at the Otowi, Albuquerque (ABQ), and San Acacia gage and the NRCS spring flow forecast at Otowi from 1997 to 2024.

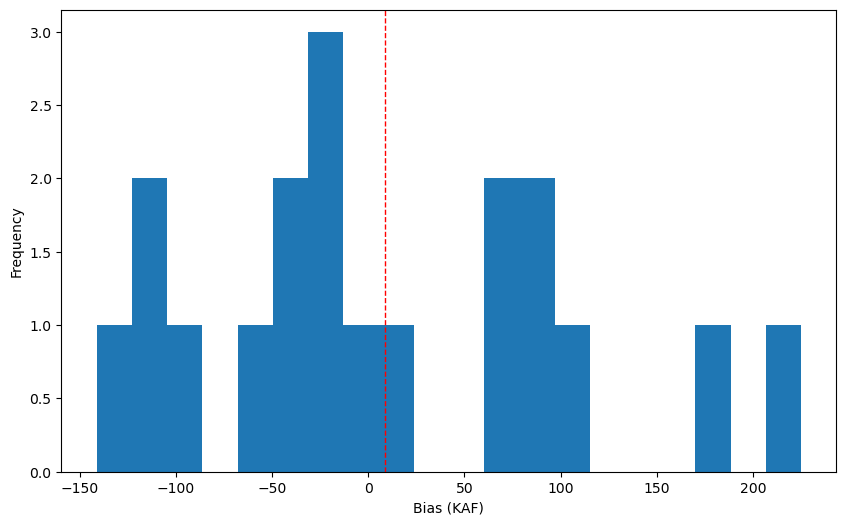

Figure S3. Histogram of the bias of the NRCS 50% exceedance forecast at Otowi in thousand acre-feet (KAF). The red line signifies the average of the biases.

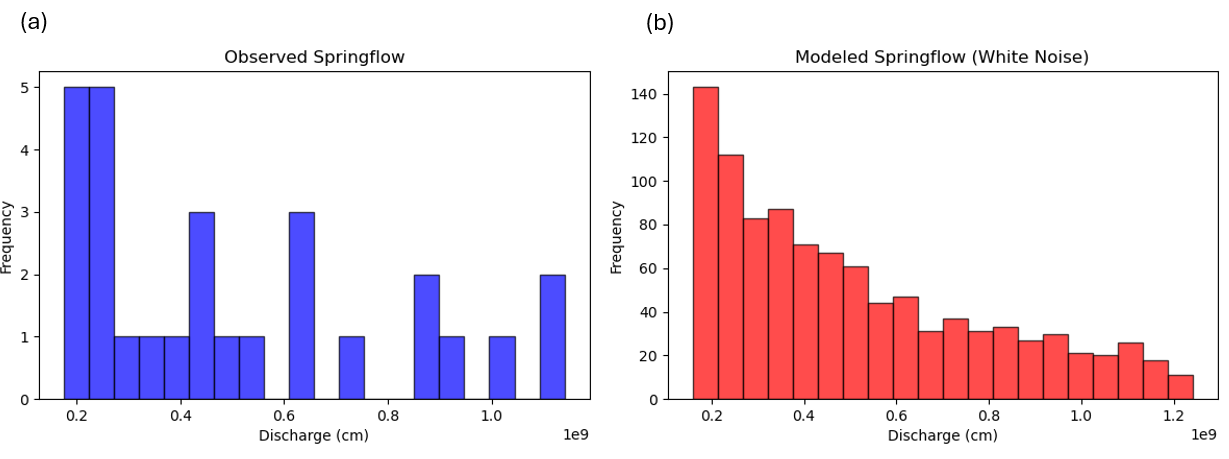

Figure S4. Observed spring flow volume from the Albuquerque stream gage (1997 – 2024) (a) and modeled spring flow (1000 samples) for Angostura reach (b) in cubic meters.

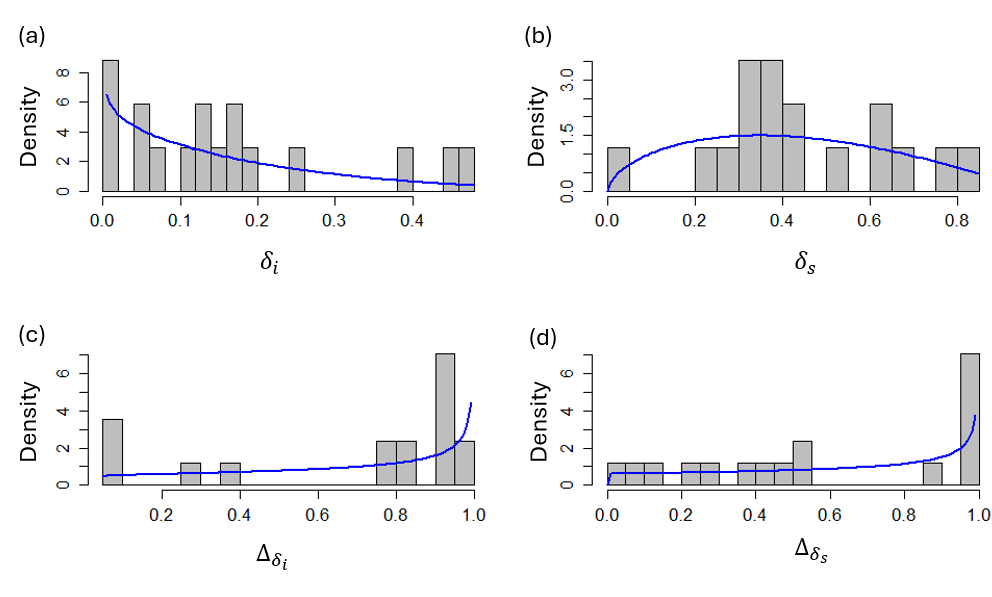

Figure S5. Proportion of river dried at least once ($\delta$) between April 1^st^ and October 31^st^, and the percentage of that drying that happened between July 1^st^ and October 31^st^ ($\Delta_{\delta}$) for Isleta (a,c; index $i$) and San Acacia (b,d; index $s$) reaches. Blue lines indicate the modeled density from the fitted beta distribution.

**Appendix D. Larval carrying capacity index prediction**

We model the larval carrying capacity index ($L$) as a function of mean annual spring flow for each reach using the general additive model. Larval carrying capacity was calculated using the daily hydrograph from the Albuquerque and San Acacia gages for the respective reaches as defined in Yackulic et al. (2022). When simulating, the larval carrying capacity index is sampled for a given spring flow at a reach by adding a normal error to the predicted value.
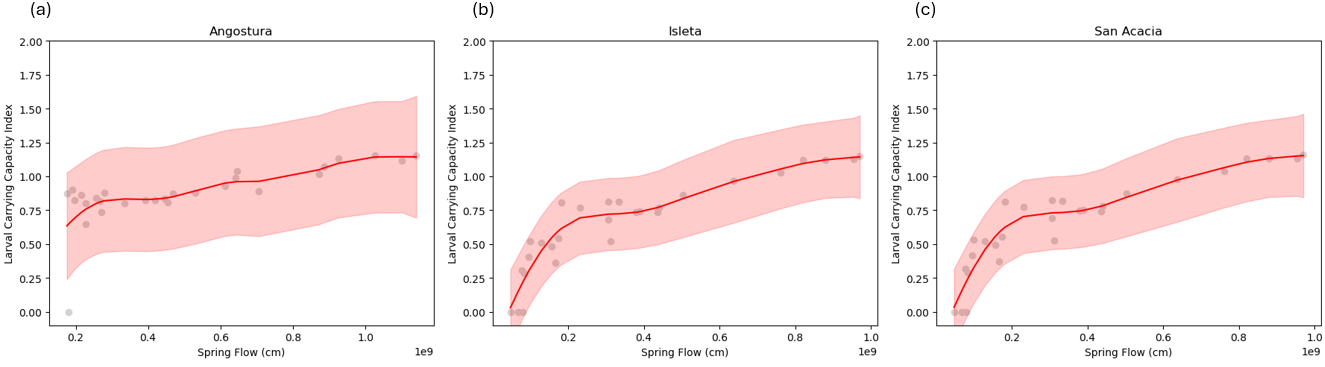

Figure S6. Relationship between spring flow and larval carrying capacity index (L) for the (a) Angostura, (b) Isleta, and (c) San Acacia reaches. Points represent observed values derived from daily hydrographs at the Albuquerque and San Acacia gages from 2002 to 2018. Red lines show the fitted relationships from generalized additive models (GAMs), and shaded ribbons indicate 95% confidence intervals.

**Appendix E. Elicitation of local quasi-extinction threshold**

Quasi-extinction thresholds are extremely hard to measure and were not estimated through the IPM in our application. To estimate this value, we held an expert elicitation session using the IDEA protocol (Hemming et al., 2018) with eight experts regarding RGSM biology. We used the four-point elicitation method from (Speirs-Bridge et al., 2010), asking the following questions to elicit what spring density (fish/km) corresponds to the quasi-extinction threshold.

1. Realistically, what do you think the lowest critical spring density threshold could be?
2. Realistically, what do you think the highest critical spring density threshold could be?
3. Realistically, what is your most likely estimate of the critical spring density threshold?
4. How confident are you that your interval, from lowest to highest, could capture the true critical spring density threshold?

The set of questions was asked in three rounds. At the end of each round, the answers were graphically displayed for the panel anonymously and were discussed to understand different interpretations and reasoning, and to examine evidence. We provided the population density estimates for each reach derived from the population size estimates from the IPM (Yackulic et al., 2022) to the expert panel, as a potential benchmark the upper boundary for the quasi-extinction threshold density.

We used the answers from the final round to derive our estimation for the quasi-extinction density threshold, $d_{ext}$. In deriving the estimation, we used each expert's lower, most likely, and upper estimate as parameters of a triangular distribution (Camus et al., 2022). Then, we picked a random expert and then sampled from its triangular distribution over 4,000 times, a number identical to the posterior distribution samples from the IPM. This method is numerically equivalent to the equal-weight linear opinion pool method to aggregate expert opinion (Howerton et al., 2023). Figure S7 shows the resulting distribution of the estimate on the quasi-extinction density threshold.

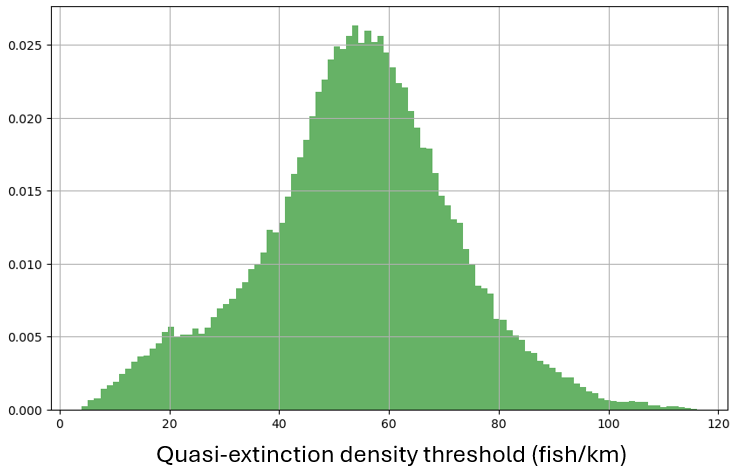

Figure S7. Distribution of quasi-extinction density threshold derived from the expert elicitation session.

**Appendix F. Deep reinforcement learning**

We used the TD3 algorithm (Fujimoto et al., 2018) to train our policy because it supports continuous action spaces, allowing us to model stocking decisions as proportions constrained to a 2-simplex (three non-negative proportions that sum to one). It also employs several techniques to stabilize training, such as target policy smoothing, delayed policy updates, and usage of multiple critic networks to mitigate overestimation bias. The algorithm uses five neural networks total: one policy network (actor), two value function networks (critic), and three target networks for each network. The actor network maps observed system states to continuous management actions, representing a deterministic policy that directly outputs production and stocking decisions. The critic networks approximate the action–value function (Q-function), evaluating the long-term expected return associated with a given state–action pair under the current policy, and thereby provide the learning signal used to update the actor. The architecture of the actor and critic networks is displayed in Figure S8. Target networks are used to compute stable temporal-difference targets for critic updates, and are themselves updated via Polyak averaging from the main networks. Two critics are used so that, when computing the temporal-difference target, the algorithm can take the minimum of the two Q-value estimates, thereby reducing the positive bias that typically arises from function approximation in Q-learning (Mnih et al., 2015). Hyperparameter values for training are outlined in Table S2.

We made small modifications to the TD3 algorithm to make the training more stable for our application. First, we transformed the actions in the sampled transitions to effective actions before inputting them into the critic network. A transition is a tuple, $(o_{t},a_{t},R_{t+1}, o_{t+1})$, that describes what action the agent took ($a_{t}$) given the observation ($o_{t}$), followed by the resulting reward ($R_{t+1}$) and the next observation ($o_{t+1}$) in a single step of the simulation. Effective actions zeroed the stocking distribution actions in spring season transitions and production actions in fall transitions to avoid training on spurious gradients produced from actions that are not relevant in respective seasons. Second, we added the exploration noise and target policy output smoothing noise ($\xi$) to only the relevant actions in respective seasons as well. This kept the agent from exploring in action dimensions that are ignored by the environment on that step.

In a training session, at every few steps, random samples of transitions were selected to update the parameters in the policy and value function networks at a small rate. In early stages of a training session, an agent deviates from the actions output by the policy using the Ornstein–Uhlenbeck (OU) stochastic process to explore the state space and encourage visitation of diverse trajectories. At later episodes of a session, the magnitude of the exploration noise is gradually reduced, allowing the agent to rely more heavily on the learned policy. This transition from exploration-driven behavior to exploitation of the optimized policy enables the agent to converge to stable strategies that maximize long-term expected rewards.

Most of the hyperparameters were tuned through a trial-and-error method, except for batch size and critic hidden layer network size. Batch size was selected via grid search over 32, 64, 128, and 256, and the number of hidden layer nodes was selected via grid search over 32, 64, 100, 128, and 256 as well.

When storing the transitions into a replay buffer when sampling for training, we standardized the observation variables to the pre-calculated mean and standard deviations for each variable. We calculated the mean and standard deviation of the variables with the observation variables collected for 1000 episodes using the current strategy. Using this fixed mean and standard deviation for standardization worked better than using running mean and standard deviation for training.

**Algorithm 1. TD3 training algorithm (Adapted from Fujimoto et al. 2018)**

**(notations here are independent from the notations in the simulation)**

Input:

Actor $\pi_{\theta}\left( o \right):outputs logits, z\in\mathbb{R}^{4}$

Simplex transform: $a=\mathrm{softmax}(z)$

Critics $Q_{\phi_{1}}\left( o,a \right), Q_{\phi_{2}}\left( o,a \right)$

Target networks: $\theta^{'},\phi_{1}',\phi_{2}'$

Replay buffer $D$

Initialize OU state $\epsilon_{t}\in\mathbb{R}^{4}$, $\sigma_{ou}=\sigma_{ou,0}$

for $e=1\ldots E$

for $t=1\ldots T$ do

Get state $s_{t}$

$\epsilon_{t}\leftarrow\epsilon_{t-1}+\theta_{ou}\left( \mu_{ou}-\epsilon_{t-1} \right)+\sigma_{ou}\xi_{t}, \xi_{t}\sim N(0,I)$

$a_{t}=softmax(\pi_{\theta}\left( o_{t} \right)+\epsilon_{t})$

Execute $a_{t}$, observe $R_{t}$, $s_{t+1}$

Store $\left( o_{t}, a_{t}, R_{t}, o_{t+1} \right)$ in $D$

Sample minibatch of N transitions $\left( s_{i}, a_{i}, R_{i}, {s^{'}}_{i} \right)_{i=1}^{N}$ from $D$

$a_{i}^{'}=\mathrm{softmax}\left( \pi_{\theta^{'}}\left( o_{i}^{'} \right)+clip\left( \xi_{i}, -c,c \right) \right),$ where $\xi_{i}\sim N\left( 0,\sigma_{c}^{2}I \right)$

Transform $a_{i},a_{i}^{'}$ into effective actions, $a_{i,eff},a_{i,eff}^{'}$

$y_{i}=r_{i}+\gamma\min\left\{ Q_{\phi_{1}^{'}}\left( o_{i}^{'},a_{i,eff}^{'} \right),Q_{\phi_{2}^{'}}\left( o_{i}^{'},a_{i,eff}^{'} \right) \right\}$

Update critics

$\phi_{1}\leftarrow argmin_{\phi} N^{-1}\Sigma_{i}\left( Q_{\phi}\left( o_{i},a_{i,eff} \right)-y_{i} \right)^{2}$

$\phi_{2}\leftarrow argmin_{\phi} N^{-1}\Sigma_{i}\left( Q_{\phi}\left( o_{i},a_{i,eff} \right)-y_{i} \right)^{2}$

If $t \mathrm{mod} d$ then

Update $\theta$ by policy gradient:

$\theta\leftarrow\theta+\alpha_{\pi}N^{-1}\nabla_{\theta}Q_{\phi_{1}}(o_{i},{softmax(\pi}_{\theta}\left( o_{i} \right)))$

Polyak averaging of targets:

$\theta^{'}\leftarrow\tau\theta+\left( 1-\tau\right)\theta^{'}$

$\phi_{i}^{'}\leftarrow\tau\phi_{i}+\left( 1-\tau\right)\phi_{i}^{'}$

end if

end for

Decay exploration noise: $\sigma_{ou}=\sigma_{ou,0}-(\sigma_{ou,0}-\sigma_{ou,E})e/E$

end for

Table S2. Hyperparameter settings for training the TD3 algorithm. Notations here are independent of the notations in the simulation model.

| Hyperparameter | Notation | Value |
| --- | --- | --- |
| actor learning rate | $\alpha_{\pi}$ | 0.0003 |
| critic learning rate | $\alpha_{Q}$ | 0.0003 |
| OU-process mean | $\mu_{ou}$ | 0 |
| OU-process mean-reversion rate | $\theta_{ou}$ | 0.15 |
| OU-process volatility initial value | $\sigma_{ou,0}$ | 0.2 |
| OU-process volatility last value | $\sigma_{ou,E}$ | 0.05 |
| buffer size | $n_{B}$ | 1,000,000 |
| batch size | $B$ | 128 |
| target update rate | $\tau$ | 0.1 |
| policy update delay | $d$ | 2 |
| target noise | $\sigma_{c}$ | 0.05 |
| target noise clip | c | 0.25 |
| maximum steps per episode | $T$ | 50 |
| number of episodes | $E$ | 6,000 |

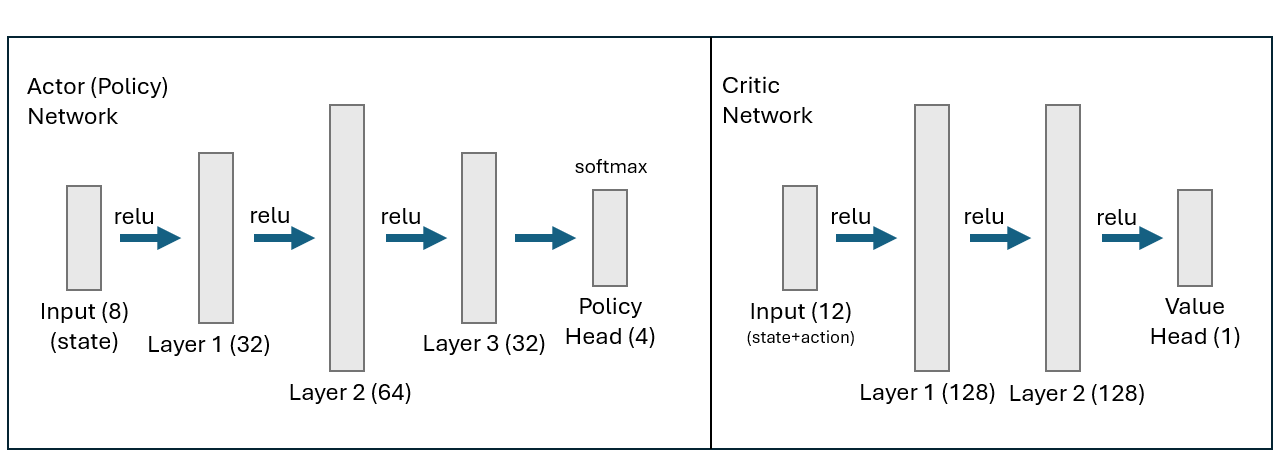

Figure S8. Network architectures for the actor (policy) and critic. Both networks are fully connected multilayer perceptrons (MLPs). Numbers in parentheses indicate the number of units in each layer.

**Reference**

Archdeacon, T. P. (2022). *Rio Grande Silvery Minnow Annual Augmentation Plan 2023-2028*. https://doi.org/10.13140/RG.2.2.29268.88968

Caldwell, C. A., Falco, H., Knight, W., Ulibarri, M., & Gould, W. R. (2019). Reproductive Potential of Captive Rio Grande Silvery Minnow. *North American Journal of Aquaculture*, *81*(1), 47–54. https://doi.org/10.1002/naaq.10068

Camus, E. B., Rhodes, J. R., Mcalpine, C. A., Lunney, D., Callaghan, J., Goldingay, R., Brace, A., Hall, M., Hetherington, S. B., Hopkins, M., Druzdzel, M. J., & Mayfield, H. J. (2022). Using expert elicitation to identify effective combinations of management actions for koala conservation in different regional landscapes. *Wildlife Research*, *50*(7), 537–551. https://doi.org/10.1071/WR22038

Fujimoto, S., Van Hoof, H., & Meger, D. (2018). Addressing Function Approximation Error in Actor-Critic Methods. *35th International Conference on Machine Learning, ICML 2018*, *4*, 2587–2601.

Hemming, V., Burgman, M. A., Hanea, A. M., McBride, M. F., & Wintle, B. C. (2018). A practical guide to structured expert elicitation using the IDEA protocol. *Methods in Ecology and Evolution*, *9*(1), 169–180. https://doi.org/10.1111/2041-210X.12857

Howerton, E., Runge, M. C., Bogich, T. L., Borchering, R. K., Inamine, H., Lessler, J., Mullany, L. C., Probert, W. J. M., Smith, C. P., Truelove, S., Viboud, C., & Shea, K. (2023). Context-dependent representation of within- and between-model uncertainty: Aggregating probabilistic predictions in infectious disease epidemiology. *Journal of the Royal Society Interface*, *20*(198). https://doi.org/10.1098/rsif.2022.0659

McKenna, C. (2019). *River Eyes Monitoring Report*.

Mnih, V., Kavukcuoglu, K., Silver, D., Rusu, A. A., Veness, J., Bellemare, M. G., Graves, A., Riedmiller, M., Fidjeland, A. K., Ostrovski, G., Petersen, S., Beattie, C., Sadik, A., Antonoglou, I., King, H., Kumaran, D., Wierstra, D., Legg, S., & Hassabis, D. (2015). Human-level control through deep reinforcement learning. *Nature*, *518*(7540), 529–533. https://doi.org/10.1038/nature14236

Myhre, A. M., Engen, S., & Sæther, B. E. (2016). Effective size of density-dependent populations in fluctuating environments. *Evolution*, *70*(11), 2431–2446. https://doi.org/10.1111/evo.13063

Osborne, M. J., Dowling, T. E., Scribner, K. T., & Turner, T. F. (2020). Wild at heart: Programs to diminish negative ecological and evolutionary effects of conservation hatcheries. *Biological Conservation*, *251*(August), 108768. https://doi.org/10.1016/j.biocon.2020.108768

Osborne, M. J., Perez, T. L., Altenbach, C. S., & Turner, T. F. (2013). Genetic analysis of captive spawning strategies for the endangered rio grande silvery minnow. *Journal of Heredity*, *104*(3), 437–446. https://doi.org/10.1093/jhered/est013

Ryman, N., & Laikre, L. (1991). Effects of Supportive Breeding on the Genetically Effective Population Size. *Conservation Biology*, *5*(3), 325–329. https://doi.org/10.1111/j.1523-1739.1991.tb00144.x

Speirs-Bridge, A., Fidler, F., McBride, M., Flander, L., Cumming, G., & Burgman, M. (2010). Reducing overconfidence in the interval judgments of experts. *Risk Analysis*, *30*(3), 512–523. <https://doi.org/10.1111/j.1539-6924.2009.01337.x>

U.S. Geological Survey [USGS], <YEAR>, USGS water data for the Nation: U.S. Geological Survey National Water Information System database, accessed <DATE>, at https://doi.org/10.5066/F7P55KJN

Yackulic, C. B., Archdeacon, T. P., Valdez, R. A., Hobbs, M., Porter, M. D., Lusk, J., Tanner, A., Gonzales, E. J., Lee, D. Y., & Haggerty, G. M. (2022). Quantifying flow and nonflow management impacts on an endangered fish by integrating data, research, and expert opinion. *Ecosphere*, *13*(9), 1–22. https://doi.org/10.1002/ecs2.4240
