## Supplemental Information 2 for "Bridging Ecological Inference and Decision Optimization for Conservation Using Artificial Intelligence"

**Supporting Information 2**

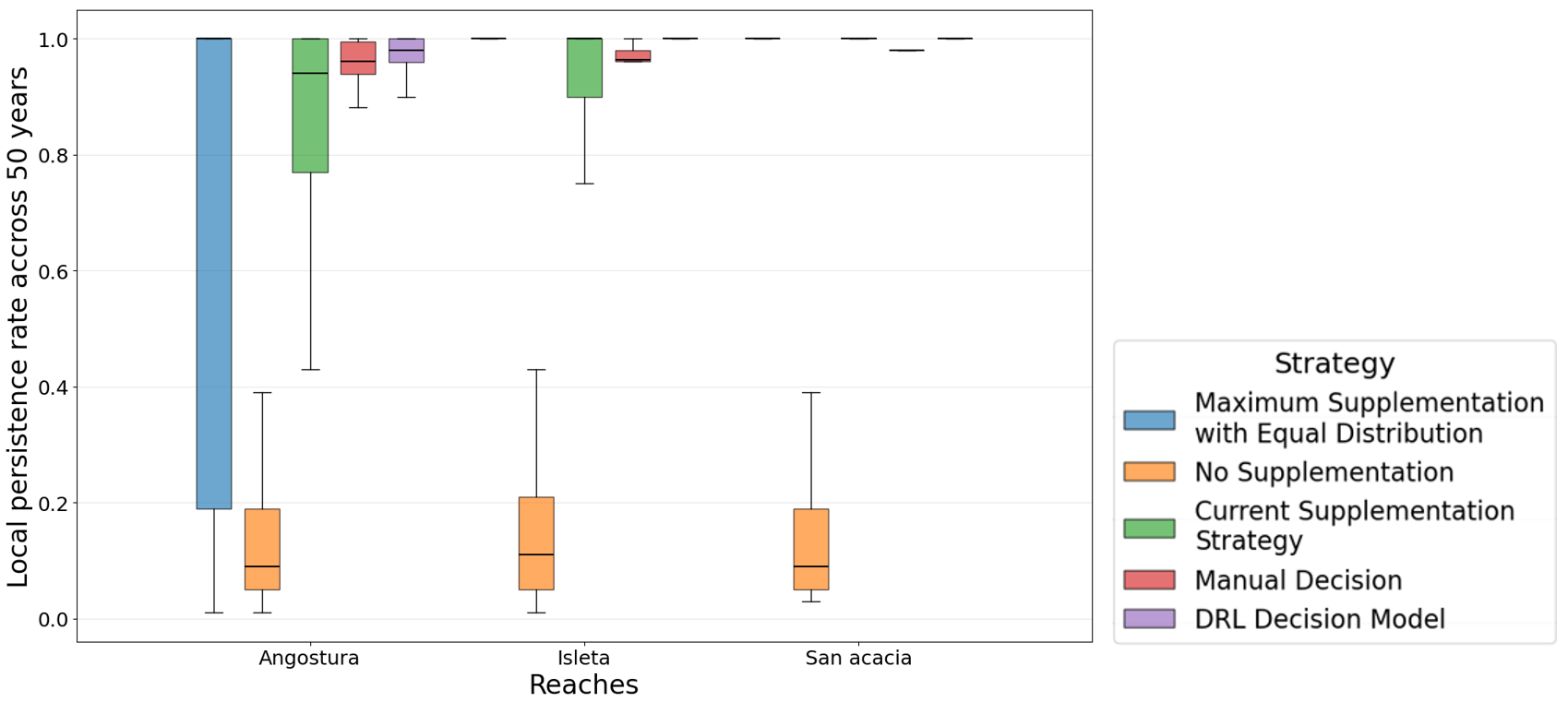

Figure S1. The local persistence rate across 50 years of supplementation for Angostura, Isleta, and San Acacia reaches using different strategies under the persistence-focused objective. Results are derived from 200 simulations.

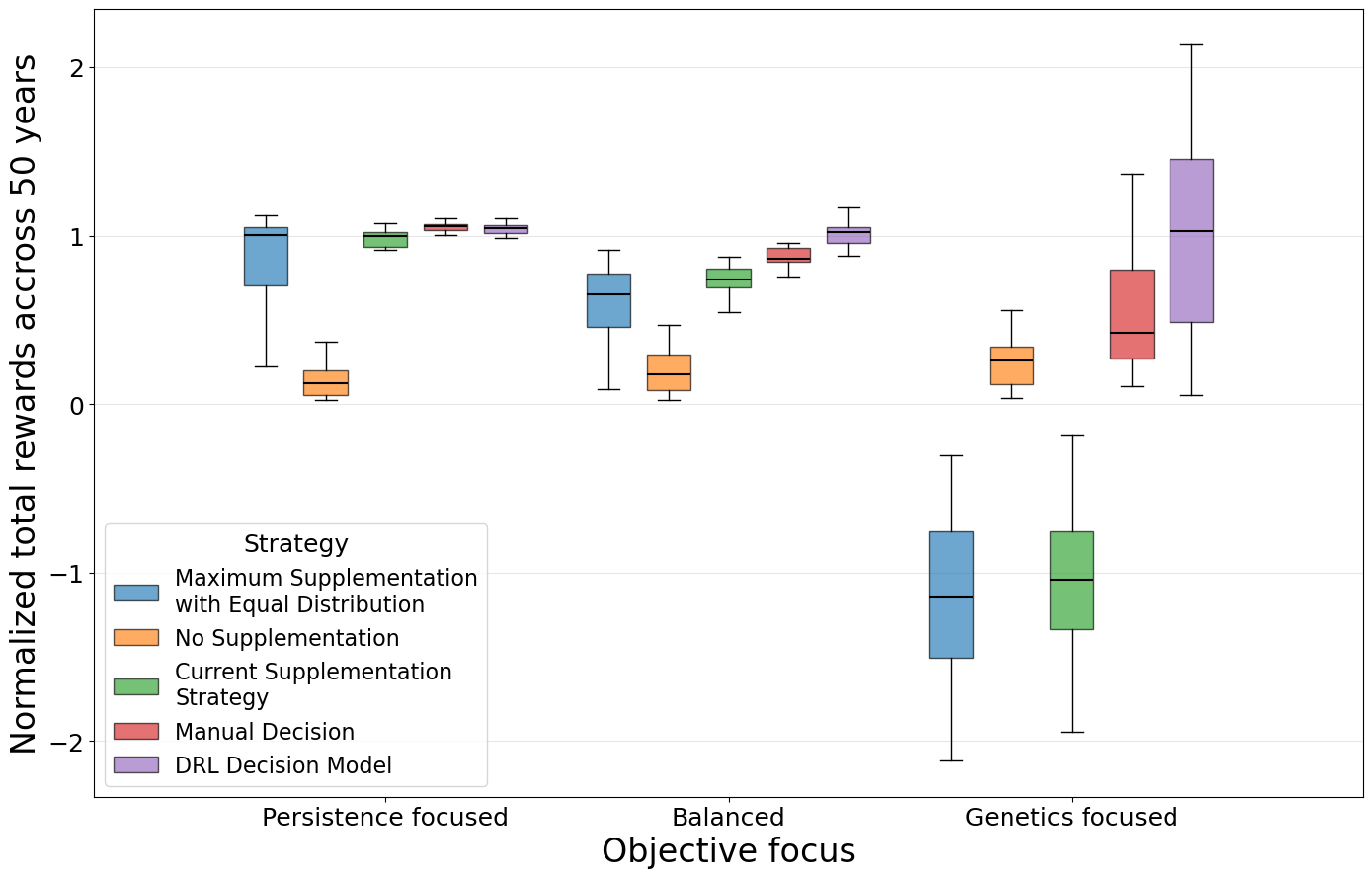

Figure S2. Average total rewards received from a 50-year supplementation using different heuristic strategies under persistence focused, balanced, and genetics focused objectives. The average total rewards are shown across 20 simulations for all strategies unlike Figure 2b in the main manuscript for equal comparison between the manual decision strategy and all the others.

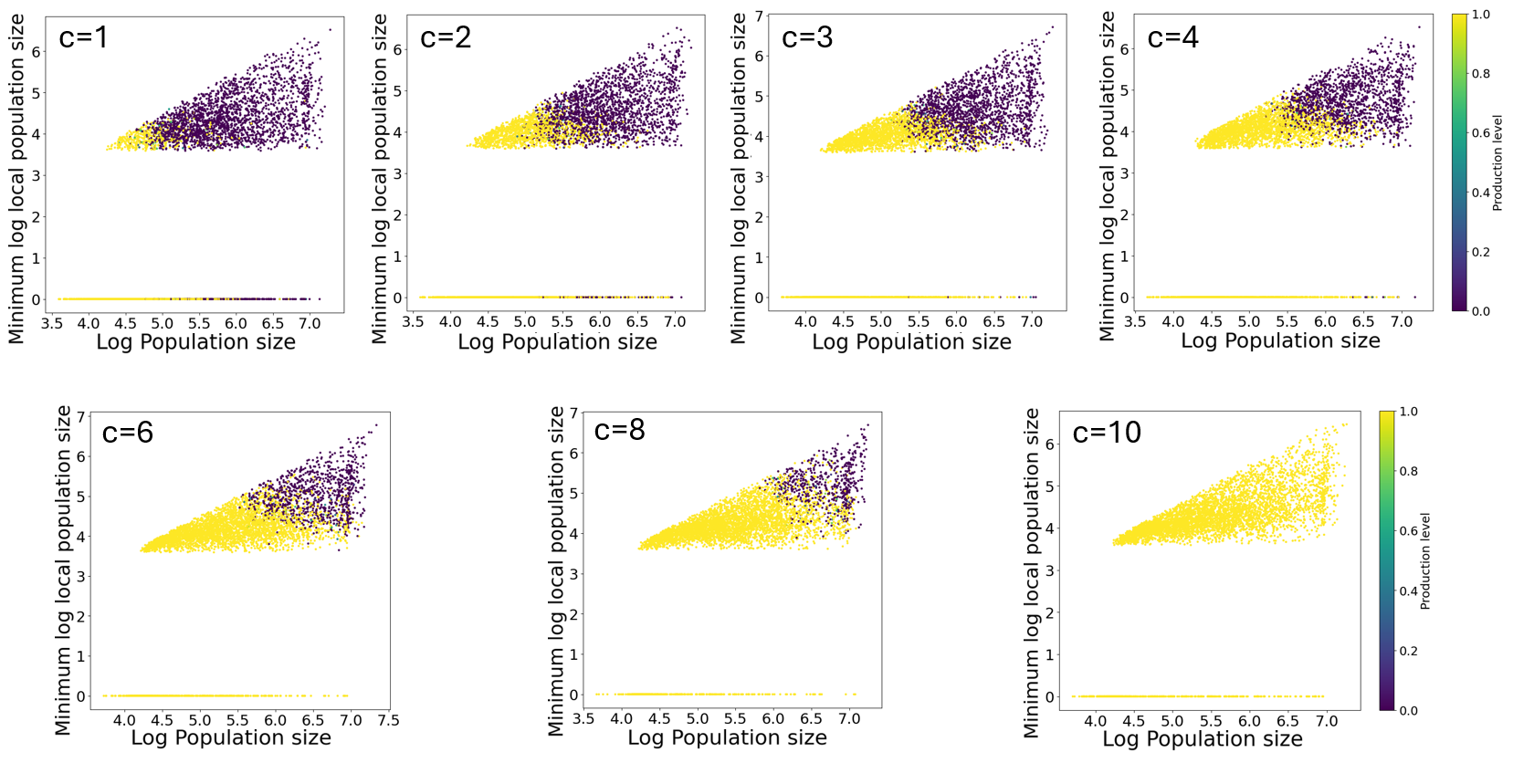

Figure S3. Annual production level decision described by log of minimum local population size and log of total population size given different value of persistence (*c*). Higher value of persistence moves the threshold of full capacity production decision up and right, producing at higher population overall. Production level of 1 equates to producing the maximum capacity of the hatchery, which is 200,000 fish. The decisions are sampled across a hundred 50-year supplementation simulations.

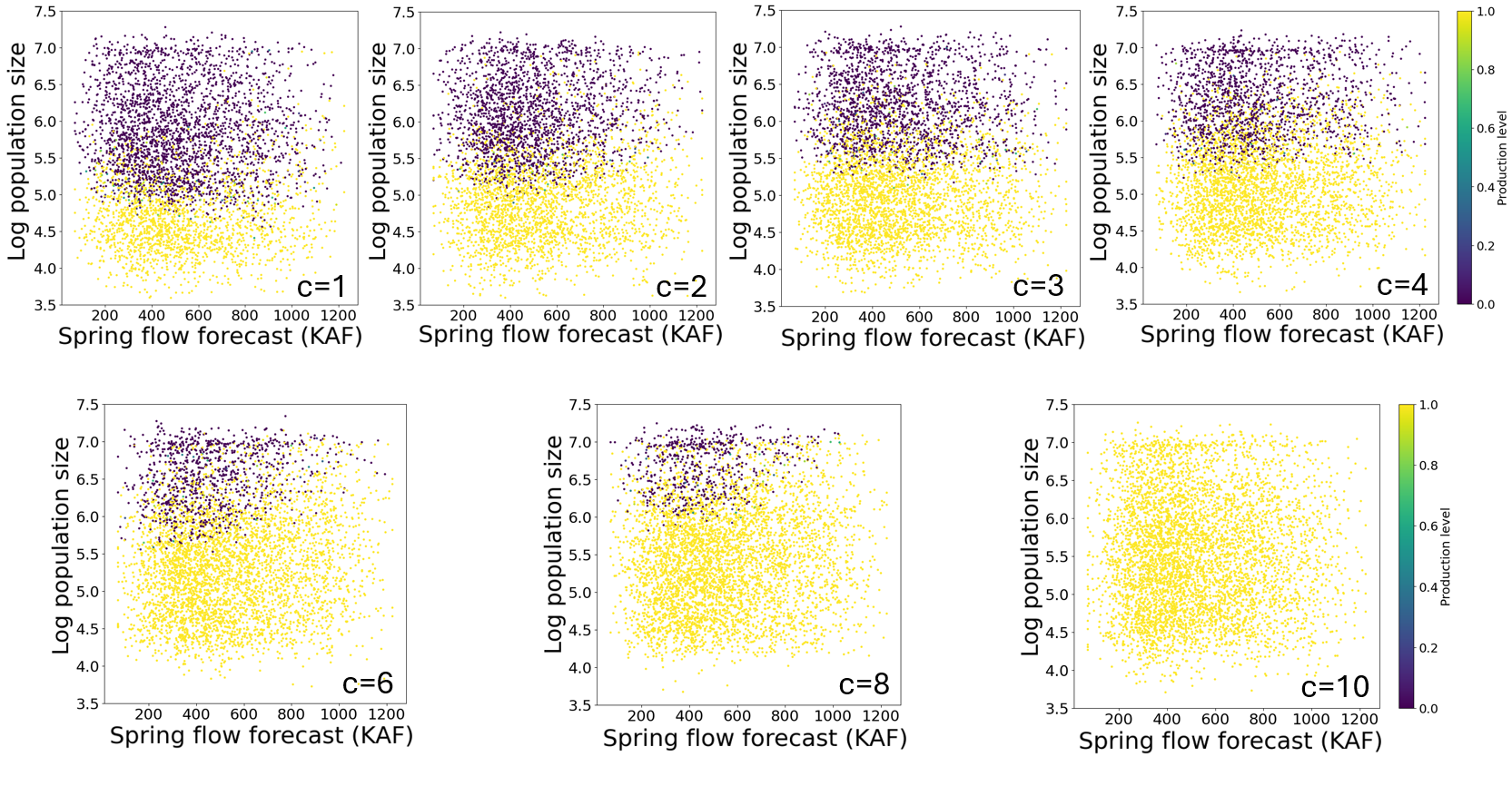

Figure S4. Annual production level decision described by log of total population size and spring flow volume forecast in thousand acre-feet (KAF) given different value of persistence (*c*). Higher value of persistence moves the threshold of no production decision to when spring flow forecast is low and population size is high, where the genetic impact per fish stocked is the greatest. Production level of 1 equates to producing the maximum capacity of the hatchery, which is 200,000 fish. The decisions are sampled across a hundred 50-year supplementation simulations.

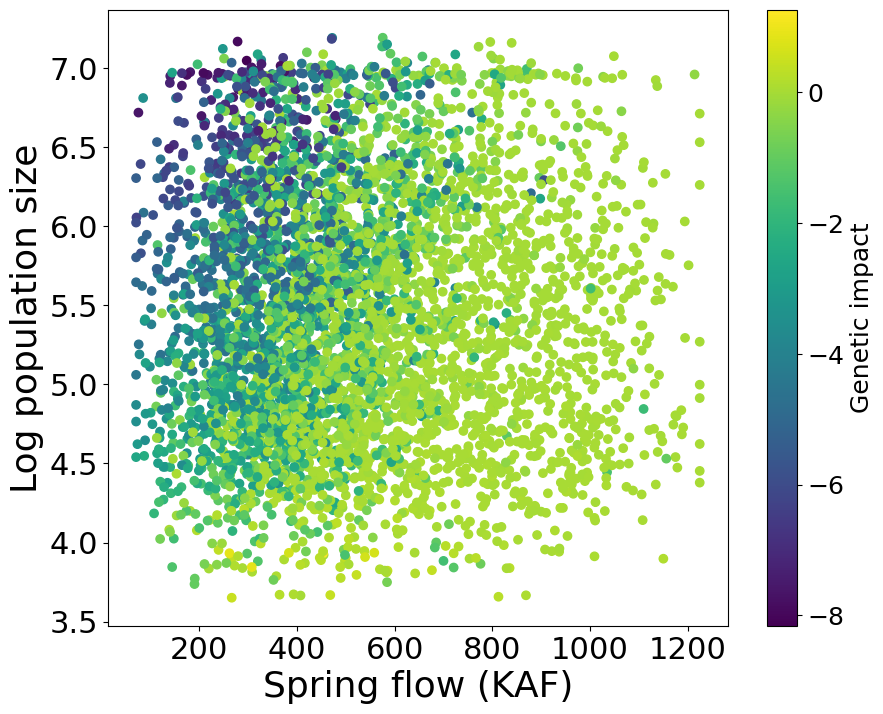

Figure S5. Genetic impact from stocking maximum capacity and distributing them equally across three local populations given the log of population size and spring flow volume in thousand acre-feet (KAF). The impacts are sampled across a hundred 50-year supplementation simulations.

Table S1. Logistic regression analysis of the annual production decision under balanced objective using covariates derived from the observed states. The area under the curve of the receiver operating characteristic curve (AUC-ROC) value is 0.98. Few annual production decisions that are between 0 and 1 were rounded to fit the logistic regression. The decisions are sampled across a hundred 50-year supplementation simulations. Covariates are standardized.

| Covariates | Coefficients | Odds Ratios | AUC drop when removed | AUC with only this variable |
| --- | --- | --- | --- | --- |
| Log_10_(total population size) | -5.1751 | 0.0057 | 0.039 | 0.943 |
| Log_10_(minimum local population size) | -3.4194 | 0.0327 | 0.009 | 0.918 |
| Coefficient of Variation between local population sizes | 1.4537 | 4.2791 | 0.004 | 0.552 |
| Spring flow | 0.5712 | 1.7704 | 0.002 | 0.533 |
| Intercept | 3.978 |  |  |  |

Table S2. Dirichlet regression analysis of the annual stocking decision under balanced objective using covariates derived from the observed states. Coefficients for the stocking proportion to Angostura reach is selected as the reference category and its summary is thus omitted. Covariates are standardized. The average congruence between the predicted and observed stocking distribution is 0.82. Congruence is defined as the proportion of the overall fish allocated to the same reaches between the predicted and observed.

| Covariates | Coefficients  for stocking proportion in Isleta | Standard Error for stocking proportion in Isleta | P-value for stocking proportion in Isleta | Coefficients  for stocking proportion in San Acacia | Standard Error for stocking proportion in San Acacia | P-value for stocking proportion in San Acacia |
| --- | --- | --- | --- | --- | --- | --- |
| Log_10_(Angostura population size) | 0.27869 | 0.016 | < 2e-16 | 0.06163 | 0.01559 | 7.72E-05 |
| Log_10_(Isleta population size) | -0.83245 | 0.0232 | < 2e-16 | -0.36066 | 0.02878 | < 2e-16 |
| Log_10_(San Acacia population size) | -0.12859 | 0.02801 | 4.41e-06 | -0.59965 | 0.02735 | < 2e-16 |
| Proportion of total population in Angostura | 0.21283 | 0.09843 | 0.0306 | 0.66366 | 0.09497 | 2.78E-12 |
| Proportion of total population in Isleta | -1.11088 | 0.10605 | < 2e-16 | 0.3606 | 0.10806 | 0.000847 |
| Intercept | 2.48859 | 0.11939 | < 2e-16 | 2.36981 | 0.12693 | < 2e-16 |
